# Prevalence, causes and impact of *TP53*-loss phenocopying events in human tumors

**DOI:** 10.1101/2022.11.01.514743

**Authors:** Bruno Fito-Lopez, Marina Salvadores, Miguel-Martin Alvarez, Fran Supek

## Abstract

*TP53* is a master tumor suppressor gene, mutated in approximately half of all human cancers. Given the many regulatory roles of the corresponding p53 protein, it is possible to infer loss of p53 activity -- which may occur from trans-acting alterations -- from gene expression patterns. We apply this approach to transcriptomes of ~8,000 tumors and ~1,000 cell lines, estimating that 12% and 8% of tumors and cancer cell lines phenocopy *TP53* loss: they are likely deficient in the activity of the p53 pathway, while not bearing obvious *TP53* inactivating mutations. While some of these are explained by amplifications in the known phenocopying genes *MDM2, MDM4* and *PPM1D*, others are not. An analysis of cancer genomic scores jointly with CRISPR/RNAi genetic screening data identified an additional *TP53*-loss phenocopying gene, *USP28*. Deletions in *USP28* are associated with a *TP53* functional impairment in 2.9-7.6% of breast, bladder, lung, liver and stomach tumors, and are comparable to *MDM4* amplifications in terms of effect size. Additionally, in the known CNA segments harboring *MDM2*, we identify an additional co-amplified gene (*CNOT2*) that may cooperatively boost the *TP53* functional inactivation effect. An analysis using the phenocopy scores suggests that *TP53* (in)activity commonly modulates associations between anticancer drug effects and relevant genetic markers, such as *PIK3CA* and *PTEN* mutations, and should thus be considered as a relevant interacting factor in personalized medicine studies. As a resource, we provide the drug-marker associations that differ depending on *TP53* functional status.

## Introduction

Mutations in the *TP53* tumor suppressor gene are a very common feature across almost all types of human cancer. These mutations abrogate or reduce *TP53* activity via various mechanisms: dominant-negative acting missense mutations, loss-of-function missense, nonsense, frameshift indel, splice site, or synonymous mutations, or copy number losses that frequently delete one *TP53* allele while the other allele is inactivated by a mutation. That such *TP53* genetic alterations occur at high frequency in many cancer types implies that they have very strong selective advantages for the expanding cancer cell clones (*1, 2*); indeed this is borne out in experimental data on cell lines and animal models of cancer (*3, 4*).

The large selective advantage of *TP53* losses are consistent with its roles in arresting the cell cycle or triggering apoptosis upon threats to genome integrity. *TP53*-null cells better tolerate genomic instability, which can result from endogenous causes, most prominently oncogene-overexpressing and thus replication-stress inducing cancerous genetic backgrounds. Consistently, *TP53*-mutant tumors have higher frequencies of segmental copy number alterations (CNA), whole-genome duplications, and overall mutation rates (*5, 6*). Moreover, *TP53*-null cells better tolerate DNA damaging conditions that would normally trigger cell cycle checkpoints, such as those resulting from DNA-acting drugs or radiation (*7, 8*). Consistently, *TP53*-mutation bearing tumors tend to be more resistant to various cancer chemotherapies (4, 9–11) and radiotherapy (*10–12*), and more aggressive i.e. *TP53* R273 and R248 mutants are associated with accelerated cancer progression in colorectal tumors (13).

The frequency of *TP53* mutations --highest of all cancer genes, standing at 37% in the TCGA cohort-- indicates that most cancers benefit from the loss of *TP53*. However, there are nonetheless many tumors which do not bear a mutation in *TP53*. A part of those is explained by genetic events that phenocopy *TP53* loss i.e. that have similar downstream phenotypic consequences as *TP53* loss itself. There are three established examples of *TP53* loss phenocopying events occuring in tumors. Most prominently, this is the amplification of the *MDM2* and *MDM4* oncogenes and overexpression of the corresponding proteins. These negatively regulate *TP53* protein levels by promoting its proteasomal degradation, and that otherwise inhibit *TP53* activity by binding to its transactivation domain(14–16). The third implicated gene is *PPM1D*, whose amplification overexpresses a serine/threonine phosphatase acting upon various targets including *TP53*, reducing its activity. (We note that *PPM1D* can also be affected by point mutations that result in gain-of-function(17–19))

Given the strong selective advantages of the *TP53* activity loss in cancer evolution, we hypothesized that *TP53* loss phenocopying in human cancers extends beyond these known examples of *MDM2, MDM4* and *PPM1D* alterations. If indeed other common mechanisms of *TP53* phenocopying exist, this would be relevant to predicting tumor cell response to various drugs, and to predicting tumor aggressiveness, thus having implications to personalized medicine. Because *TP53* loss has clear consequences on the mRNA expression levels of various downstream targets (*4, 21*), the *TP53*-null-like phenotype can be inferred from large scale transcriptomic data (20–23). Here, we apply a statistical framework to jointly analyse ~966 cancer cell line and ~8000 tumor genomes and transcriptomes, to identify additional *TP53* phenocopying genetic events and impact on drug sensitivity. We find that TP53 loss phenocopies are remarkably common across tumors and cancer cell lines, and we identify *USP28* deletions as one cause of *TP53* loss phenocopying, and reveal many links between drugs and their targets that are modulated by *TP53* activity.

## Results

### Inferring the functional *TP53* status of tumors from transcriptomes

We developed a machine learning method to detect *TP53* phenocopies in tumors and cell lines, integrating RNA-seq data with *TP53* mutation data in a logistic regression, regularized with an Elastic Net penalty (very similar cross-validation accuracy was obtained with Ridge or Lasso penalties; see Methods). Regression models were trained using cross-validation on mRNA levels of ~8000 tumor samples from the TCGA project, across 20 different cancer types, controlling for cancer type. In addition to using this global analysis mRNA expression levels to infer the functional *TP53* status state of each tumor, we also identified the expression patterns of which genes are associated with *TP53* status. Tumors with *TP53* putatively causal mutations were included as a positive set (*TP53* status was categorized according to GDSC methodology; see Methods). Previously known phenocopying events (*MDM2, MDM4* and *PPM1D* amplifications), as well as samples with *TP53* deletions were excluded from the training set (these known phenocopying events will be used to calibrate decision thresholds; see below). Our classifier learned a combination of relevant gene weights that differentiate samples with an aberrant *TP53* activity. Tumor samples that are not *TP53* mutated (by GDSC criteria), but are classified as mutated by the machine learning model are considered to be *TP53* phenocopies.

Our classifier showed a high performance with an area under the receiver operating characteristic (AUROC) curve of 96% in cross-validation on TCGA tumors (out-of-sample accuracy), and 95% on the testing set (consisting of 10% of the samples held out from training set, Fig.1A). Thus, we were able to often correctly detect *TP53* status in unseen tumor samples the classifier was not exposed to, with an area under precision-recall curve=0.9654. The *TP53* loss phenocopy scores for each TCGA tumor sample and the cancer cell lines are provided in Supplementary Data 1.

**Figure 1.**
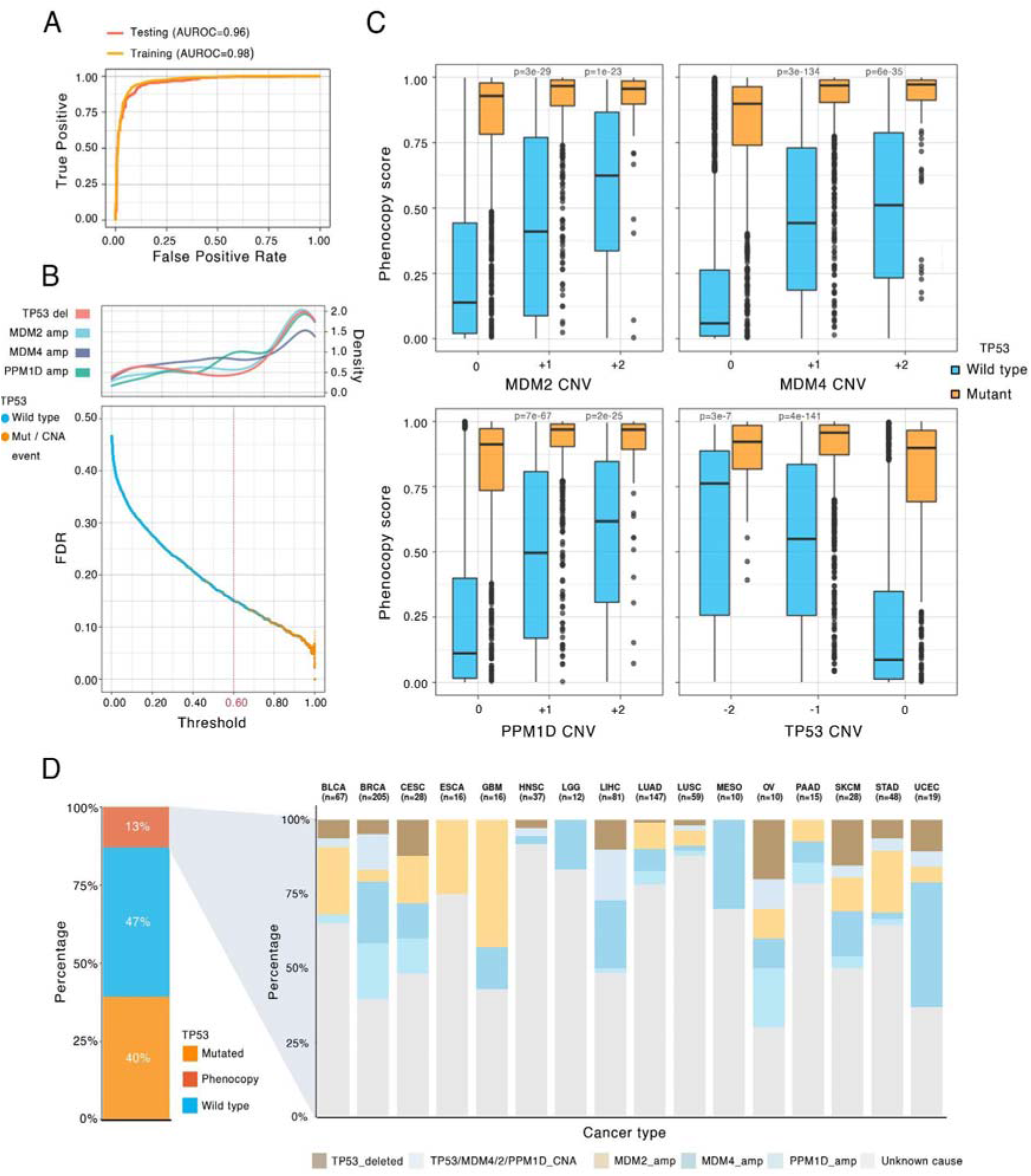
Evaluation of the functional *TP53-loss* score classifier and prevalence of *TP53* loss phenocopying events in cancer. A. Receiver operating characteristic (ROC) curve and area under the ROC (AUROC) curve for training and testing sets in TCGA tumor transcriptomes. B. Bottom: False discovery rate (FDR) for each tumor sample. X axis is the classification threshold for each tumor sample. The general threshold used for classification (0.6) is highlighted. Top: the histogram of frequency of CNV events (“density” refers to smoothed relative frequency) affecting *TP53* and the known phenocopying genes *MDM4, MDM2* and *PPM1D* at various phenocopy-score thresholds. C. *TP53* loss phenocopying score stratified by 3 known phenocopying CNA events and by *TP53* deletions. Data points are tumor samples coloured by *TP53* status; boxes show median, Q1 and Q3, while whiskers show range (outlying examples shown as separate dots). X axis represents the GISTIC thresholded CNV of each given gene. Tumor samples with deletions in the corresponding genes (for MDM2, MDM4 and PPM1D) and amplifications (TP53) are omitted for simplicity. P values represent results from the t-test comparison of the TP53 phenocopy score between each CNV category to neutral CNV (0) category in TP53 wild-type samples. D. *TP53* functional status classification across TCGA cancers. Left: pan-cancer; “*Phenocopy’’* refers to *TP53-loss* phenocopying tumors according to the classifier in panels A, B. Right: showing only *theTP53* loss phenocopying tumor samples, stratified by cancer type and by cause of the phenocopy. Tumor samples harbouring a known event that affects *TP53* functionality are shown with colours, and the remaining *TP53*-loss phenocopy tumors are labelled as “Unknown cause

Out of the ~12000 genes available to the classifier, 217 genes were deemed relevant for *TP53* status classification (non-zero coefficients; gene score provided in Supplementary Data 2). These represent a sparse (but not necessarily exhaustive) set of genes that are, considered together, highly informative for predicting *TP53* status.

Expectedly, many of the classifier’s most relevant genes are known to be related to *TP53* functionality. For instance, *apoptosis-enhancing nuclease (AEN*) was the gene with the highest absolute importance score. This exonuclease is a direct *TP53* target whose expression is regulated by the phosphorylation of *TP53* and its tumor suppressor role has been reported (25). Tumors with a high expression of AEN are expected to be p53 functional, and indeed highly expressed *AEN* was associated with *TP53* WT status in our classifier’s coefficients. On the other extreme, *COP1*, a ubiquitin ligase that acts as an important p53 negative regulator, was the strongest coefficient associated with *TP53* mutated status in the classifier (26). We further performed a GO enrichment analysis, revealing that top functional enriched sets were related to apoptotic signals, supporting the biological rationale underlying this set (Supp Fig. 1A).

Most enriched pathways were*: Intrinsic apoptotic signalling pathway in response to DNA damage by p53 class mediator* (8.1-fold enrichment, FDR=4.2%), *Pyrimidine deoxyribonucleoside monophosphate biosynthetic process* (47.4 fold enrichment, FDR=1.9%) and *Response to UV-B* (17.2 fold enrichment, FDR=3.7%) (ShinyGO, see Methods).

Our classifier extends recent gene expression-based models for TP53 functionality (20–23) by being able to generalize across both tumor and cancer cell lines (important for identifying drug sensitivity associations, see below), and moreover it can provide calibrated FDR estimates for TP53 status of each tumor or cell line. In particular, to assess the reliability of the individual predictions from the model, FDR for each tumor was computed via the analysis of cross-validation precision-recall curves (Fig.1B). The previously known phenocopies (*MDM2, MDM4* and *PPM1D* amplifications) and *TP53* deep deletions, which were held out from the training set, were largely scored as *TP53* mutated. Tumors harbouring a known phenocopying amplification were assigned higher scores than the rest of *TP53* wild-type tumors (means=0.56 and 0.27 respectively, p=1e-65 by t-test). Cells harbouring a *TP53* deep deletion also had higher scores (mean *TP53* deleted=0.47, mean *TP53* not deleted=0.27, p=1e-08). Our choice of threshold to detect *TP53* phenocopied tumors was set based on these known phenocopies, conservatively, corresponding to score >0.6, Methods; Fig.1B).

This resulted in an empirical FDR estimated at 15% (i.e. precision of 85%), based on the known *TP53* mutations. Importantly this 15% is a conservative upper-bound estimate of FDR, since it is based on the assumption that there do not exist any unknown *TP53* phenocopying events: it classifies all high-scoring *TP53 wild-type* tumors as false positives. Conversely, using the known phenocopying events we estimate a lower-bound recall (sensitivity) of this classifier at 63% (Fig. 1B). Again, this estimate is conservatively biased, since it is not a priori known whether every copy number gain in *MDM2/MDM4/PPM1D* causes a phenocopy; some low-level gains may not have effects and thus would appear as false-negatives.

To additionally validate the classifier, we inspected the relationship between known phenocopy genes’ allele copy-number (see Methods), and the *TP53* phenocopy score. There were significant positive correlations between three known phenocopying genes copy-number, and the *TP53* phenocopy score in *TP53* wild-type tumors (Fig.1C).

The prevalence of phenocopying events was substantial: overall 12 % of all tumor samples were redefined into a *TP53* mutated-like category (Fig.1D) by our criteria. Different cancer types display different phenocopy frequencies (Fig.1D), overall frequency ranging from 19% for breast cancer (BRCA cancer type) to 3% for B-cell lymphoma (DLBC cancer type, overall phenocopy frequencies are shown in Supp Fig. 1B). For instance, most breast cancer *TP53*-phenocopied tumors derive from previously known events i.e. the *MDM4/MDM2/PPM1D* amplifications are the most common event, while a remaining 27% of the phenocopies (5% of all breast cancer samples) is not associated with a known phenocopying event (proportion shown for every cancer type Fig.1D). We do note that it is still possible that individual examples of tumor may be erroneously classified as TP53-deficient at this threshold. More generally, 51% of *TP53*-loss phenocopied tumor samples across all cancer types were not linked with one of the three known genes nor a CNA deletion in *TP53* itself, suggesting that additional *TP53* phenocopying mechanisms are commonly occurring in tumors.

### *USP28* deletion phenocopies a *TP53* mutated state in tumors

Prompted by the abundance of tumor samples that are functionally *TP53* null but lacking an obvious *TP53* loss or a known phenocopying event, we sought to identify other phenocopying genes across all cancer types. We designed a custom associationtesting methodology that combines six different statistical tests across four different genomic data types with this goal (see Methods).

In brief, our methodology is based on the rationale that genes that cause a phenocopy via altered dosage at DNA and mRNA levels should exhibit a distinct copy number variant (“CNV” tests) and also gene expression (“GE” tests) pattern. Each of these two genomic data types is considered in two tests, one comparing *TP53* phenocopying against *TP53* wild-type tumors, and other comparing *TP53* phenocopying against TP53-mutant tumors, for a total of four tests. As two additional tests, we considered external data from genetic screens across large panels of cancer cell lines (28,29). In particular we test for significant codependency scores, explaining how a knockout (“CRISPR”) or knock-down (“RNAi”) of a candidate phenocopying gene affects fitness across a panel of cell lines, when compared with the fitness profile of a *TP53* knockout/knock-down across the same panel(30, 31). An example supporting the use of this methodology that combines cancer genomic analysis and genetic screening data analysis, a CRISPR knockout of the known *TP53* negative regulator *MDM2* decreases cell line fitness, in a manner anticorrelated to a *TP53* knockout across cell lines. (Supp Fig. 3A)

In summary, we tested differences of tumor genomics CNV and GE patterns (two tests each as above), additionally considering “CRISPR” and “RNAi” test scores from genetic screens, for each gene, performing tests stratified by cancer type. Our final score combines each of the 6 tests together providing a ranking of potential *TP53* phenocopying genes.

As anticipated, top 3 prioritization scores correspond to *MDM2, MDM4* and *PPM1D* genes (Fig. 2A). Following those known *TP53* phenocopies, the gene *USP28* was the 4th ranked gene in terms of overall statistical significance (p=5.9e-07, combined across all six tests), and in particular scored highly on CRISPR codependency (pan-cancer score for *USP28*=0.54, compared with −0.71 for *MDM2* and −0.53 for *MDM4*). A break-down of our *custom prioritization* scores by different cancer types is provided in Supplementary Figure 2. We note that, in contrast to *MDM2* and *MDM4*, it is the deletions not amplifications of *USP28* that were associated with *TP53* phenocopies; this is reflected in the mirrored direction of the codependency score. *USP28* encodes a deubiquitinase enzyme with substantial evidence from previous biochemistry and cell model studies that link it to p53 activity. In particular, *USP28* was linked to DNA damage apoptotic response through the Chk2-p53-PUMA pathway (32). Recent evidence suggests that the *TP53BP1-USP28* complex might positively regulate p53 and influence arrest after centrosome loss and prolonged mitosis (33). It has been proposed that *TP53BP1-USP28* complexes could counteract *MDM2*-dependent p53 ubiquitination (34). Additional studies have linked *USP28* loss with a defective apoptotic response (35). A 10% of the total of 437 tumors classified as *TP53* loss phenocopied but with an undefined source (Supp Fig.1B) had a *USP28* deletion.

**Figure 2:**
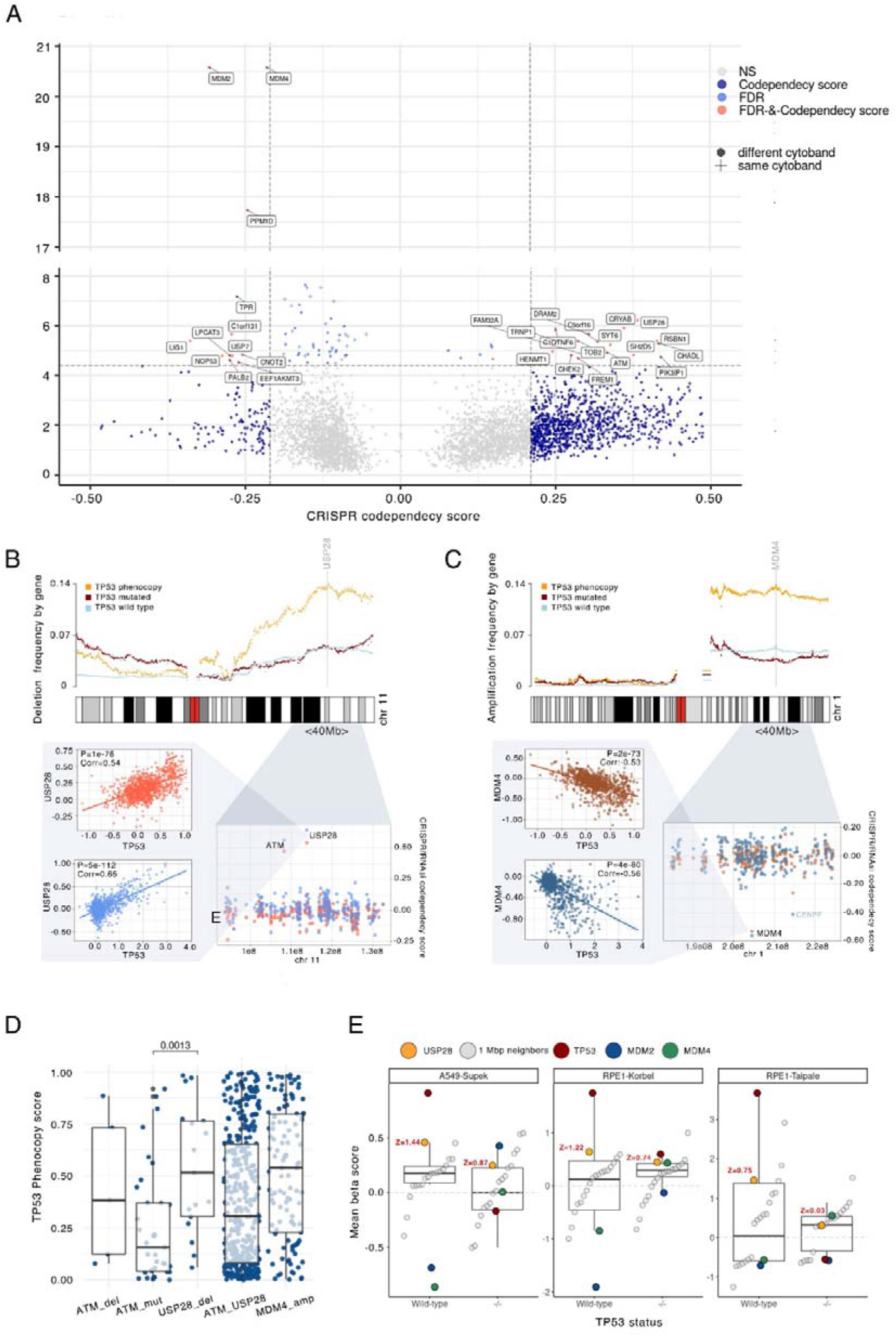
Transcriptomics scores predicting phenocopying events can pinpoint causal genes in CNA-affected chromosomal segments. **A.** Prioritization score of genes for *TP53* loss phenocopying effects. Y axis shows gene significance (FDR) when combining six statistical tests (two cancer genomic/transcriptomic, and two based on CRISPR and RNAi screens), and further pooling p-values across cancer types; see Methods for details. X axis represents the effect size from the CRISPR codependency score of each gene. Crosses represent gene neighbours (same cytoband) to a known phenocopying gene. Relevant hits in terms of FDR and codependency score are labelled. Shown thresholds for effect size and significance were determined based on scores of known phenocopy events (Methods). **B.** Top: CNV frequency in tumors, and their associations with *TP53* phenocopy transcriptomic scores, of the segment of chromosome 1 containing *MDM4*. Each dot represents one gene, while colours represent groups of tumor samples by *TP53* status. Bottom: A zoomed-in view of a commonly amplified region of the chromosome, showing the CRISPR (blue) and the RNAi (red) *TP53*-codependency scores for each gene. The determination of the *TP53* codependency score is shown for the top score of the region (left panels), showing the actual CRISPR and RNAi fitness effects for the *MDM4* disruption (Y axis) across many cell lines (dots), compared to *TP53* disruption fitness effects (X axis) across the same cell lines. **C.** Same as (B), but for *USP28*, a gene we identified to be associated with a *TP53* loss phenocopying via a deletion. Here, the y axis on the top plot shows frequency of gene deletions in tumors, divided by *TP53* functional status, whereas panel B shows frequency of amplification. Bottom plots are the same as in B. **D.** Comparison of the TP53 phenocopy score of *USP28* CNV deletions (by negative GISTIC score), *ATM* deletions, *ATM* mutations and MDM4 amplifications. Each dot represents a tumor sample. Only TP*53 wild-type* samples were considered. P-values by Mann-Whitney test. **E.** Fitness effect of *USP28* knock-out in TP53 *wild-type* and mutant isogenic cell lines. Comparison of the mean beta score (fitness effect upon CRISPR gene disruption, y-axis) of *USP28*, with the mean beta scores of genes located within its 1Mbp immediate surroundings as negative controls (“1 Mbp neighbours”, see Methods). Genes *TP53, MDM2*, and *MDM4* are also shown as a reference. x-axis bottom labels indicate the *TP53* status of the cell line. *USP28* Z-scores, comparing to the distribution of neighbouring genes, are plotted in red (see Methods)

Overall, diverse experimental evidence from genetic screens strongly supports our identification of *USP28* deletions as p53-loss phenocopying events, and our genomic analysis suggests a widespread distribution of causal *USP28* deletions across human tumors.

Additional hits from this association study might provide promising genes for follow-up. For instance, *MSI2* was the 5th most highly prioritized gene, predicted to phenocopy *TP53* loss by amplification. *MSI2* encodes a transcriptional regulator that has been recently identified as an oncogene in hematologic and solid cancers (36–38). Similar results to CRISPR analyses were observed using RNAi screening codependency scores, further supporting the role of *USP28* loss in the *TP53* phenocopying, as well as *MSI2* gains (Supp Fig. 3B). Other apoptosis-related genes such as *DRAM2, CHEK2*, or *ATM* (39–41) were also in the prioritized genes in our analysis albeit at more modest statistical significance. Of note, the *TPR* gene also had a highly significant codependency score but was driven by a single cancer type (kidney) and thus with less clear relevance to diverse tumor types.

### Phenocopy scores prioritize causal genes in CNA-affected chromosomal segments

Amplifications of certain chromosomal segments or whole arms in case of *MDM2, MDM4* and *PPM1D* commonly underlie *TP53* phenocopies. Such CNA genetic events in cancers however often affect multiple adjacent genes, where an open question in cancer genomics is which of the gene or genes in the affected segment are causal (42). We hypothesized that the known *TP53* phenocopying gene CNA segments might in some cases harbor more than one causal gene. Our combination-test approach can prioritize genes with enriched gene expression and CNA in our *TP53* phenocopying group. Considered together with CRISPR and RNAi codependency, this method provided a plausible ranking of possible *TP53* loss phenocopying genes. Applied globally, this identified *USP28* as a novel phenocopying gene (see above). To more formally study if the *USP28*-adjacent genes could contribute to this, we considered that the same method could be applied on a local scale: examining profiles of CNVs and our genomic prioritization scores would be able to single out the causal gene(s) in the chromosomal segment of recurrent CNA.

As a control for this approach, we sought to confirm previously known phenocopies. Indeed, *MDM4* amplification was found to be enriched in the *TP53*-phenocopying group of tumor samples, but not in the rest of tumor groups --the *TP53* mutant and the non-phenocopying *TP53* wild-type (Fig. 2B). The local profile of this enrichment for the chromosome 1q segment 32.1 peaks at the *MDM4* gene and falls off towards its flanking genes (Fig. 2B). Our CRISPR and RNAi data analysis, consistently, indicate *MDM4* as the gene with the strongest effect in the region (Fig. 2B). As expected, similar CNA and CRISPR/RNAi profiles were observed at *PPM1D* (Supp Fig. 3C). Next, the *MDM2* CNA enrichment score segment peak was narrower, suggesting a more focal gene amplification process (Supp Fig. 3C)

Next, we examined the shape of the local *USP28*-adjacent CNA profiles. *USP28* deletions were found to be enriched in the *TP53* phenocopying group when compared to the rest of tumor groups (2.3-fold in *TP53* w.t, 2.8-fold in *TP53* mutant). *USP28* enrichment was comparable to *MDM4* region enrichments of 2.5-3.7-fold (*TP53* wt., *TP53* mutant) (Fig. 2B, C).*TP53* phenocopying tumor samples appear to have more deletions in the *USP28* region than *TP53* wild-type (non-phenocopying) and *TP53* mutant samples. The local profile of enrichments presents a plateau-like pattern rather than a sharp peak, and *USP28* is within the top-ranked genes in the plateau however some neighbouring genes appear similarly so. Therefore, we further used the CRISPR and RNAi codependency scores to prioritize the causal genes in the segment; this score clearly distinguishes *USP28* from immediate neighbours (Fig. 2C), suggesting that *USP28* is indeed the main causal gene in the chromosomal segment.

Furthermore, this ‘local scan’ can be applied chromosome-wide, where we noted another small region on chromosome 11q.12.1-q1.13.1 modestly enriched with amplifications in *TP53*-phenocopying tumors (Supp Fig. 3D), thus raising our interest. However, neither genes in this region nor other chromosome 11 regions showed a positive CRISPR codependency score of even half of *USP28* score (Fig. 2C). We note here that the *USP28* codependency score exceeds, in absolute magnitude, the score of the known *MDM4* phenocopy (Fig. 2B, C).

In the broader neighborhood of *USP28*, the gene *ATM* seems to also be frequently deleted in the *TP53*-phenocopying tumor group, meaning *ATM* is also a candidate for the causal gene in this deletion segment at chr11 q22.3-q23.2. However, the statistical support from genomic enrichment scores (using our custom methodology for metanalysis across 6 statistical tests) for *ATM* were less strong than for *USP28* (p=1e-5 versus p=6e-7, respectively). Consistently, comparing the RNAi and CRISPR *TP53-* codependency scores of *ATM* versus *USP28* shows a stronger effect of the *USP28* knockout (*USP28* RNAi codependency score p=4.9e-112 versus *ATM* p=3e-80, in a pan-cancer analysis; Supp Fig. 3E). To further rule out that *ATM* has the causal role in this deleted segment, we considered the cases of tumors where *ATM* is disrupted by a point mutation; unlike CNA in the *ATM* gene, these cases are not commonly linked with disruptions in USP28. The *ATM* mutated but *USP28* wild-type tumors had considerably weaker TP53 phenocopy transcriptomic scores (median=0.36) than the *USP28* deleted but *ATM* non-mutated tumors (median=0.84; p=0.0013 by Mann-Whitney test; Fig. 2D). The cases where both *USP28* and *ATM* were disrupted, by deletion or mutation, had very similar phenocopy scores (median=0.73) as the *USP28* deleted but *ATM* non-mutated cases. This analysis of *ATM* mutations supports that *USP28* deletion, rather than *ATM* disruption, is the causal change in the deleted segment at chr11 q22-q23.

To validate the *USP28* finding, we analyzed an independent CRISPR data set, consisting of 3 genome-wide screens performed on *TP53* wild-type and *TP53* -/- isogenic pairs of cell lines: one on the A549 cell line pair and two on the RPE1 cell line pairs (see details in Methods). In the *TP53* wild-type background, the *TP53* k.o. increases cell fitness (as expected for a high-effect tumor suppressor gene; Fig. 2E). Thus, if the *USP28* loss were to phenocopy *TP53* loss, the *USP28* k.o. by CRISPR should also increase fitness. Indeed, it does so: compared to the 10 neighboring control genes residing within 1 Mb of *USP28*, the *USP28* k.o. has a stronger fitness effect (beta score from MAGeCK tool, see Methods) for 10 out of 10 genes in 2 out of 3 screens, and 8 out of 10 neighboring genes in the remaining screen (Fig. 2E). For *ATM*, this effect is less pronounced (Supp Fig. 3F). In 3 out of 3 cell screening experiments, *USP28* fitness effect was stronger than ATM effect (1.4-fold, 2.4-fold and 2.6-fold increased beta score). To further support this finding, we asked if the fitness gain resulting from *USP28* loss is because of downstream effects on *TP53* activity. We thus considered the isogenic cells where *TP53-/-* was ablated, in which indeed the fitness gain from *USP28* k.o. was attenuated or disappeared (Fig. 2E) compared to TP53 wild-type cells. In 2 out of 3 cell line screens, the fitness attenuation effect of TP53-/- background cells was stronger in *USP28* than in the neighboring *ATM* gene, supporting the causal role of *USP28* in that segment (Supplementary Data 3). Of note, in this analysis the effect sizes of *USP28* k.o. were less than of full *TP53* k.o., however they were still substantial: in 2 out of 3 screens considered, the fitness gain effect of *USP28* disruption was comparable in size to the fitness loss effect of *MDM4* disruption (Fig. 2E).

Overall, these analyses highlight *USP28* as the likely causal gene for *TP53* loss phenocopying via deletion CNVs in the chr11 q22-q23 segment.

### Cancer type specificity of *TP53* phenocopying events

As stated above, not every cancer type seems to be affected by the same types of phenocopies. For instance, *MDM2* amplification phenocopy occurs often in BRCA, CESC, BLCA, LUAD and STAD but it does not in HNSC, OV, MESO nor LIHC (Fig.1D). To further elucidate the tissue-specificity of *USP28* phenocopying events, we considered the prioritization scores separately for different cancer types (Supp Fig. 2). We observed that BRCA, BLCA and LUAD were the cancer types which showed the strongest signal in our prioritization score for *USP28* phenocopies, with a suggestive signal in STAD.

To elucidate the cancer type spectrum of the *USP28* phenocopies, we considered the *USP28* amplifications as a negative control (deletions are expected to phenocopy). In particular, we determined in which tumor types *USP28* deletions had a higher *TP53* phenocopy score than USP28 copy number amplified samples. As expected, statistical significance when comparing the *TP53* phenocopy score of *USP28* copy number-neutral tumor samples versus those bearing deletions was higher than comparing neutral to amplifications. This supports that the impact of *USP28* deletions on *TP53* loss phenocopy score was stronger than for the amplification CNVs. The strongest effect was found in BLCA, STAD, BRCA, LIHC and LUAD (Fig. 2E). In further support of this tissue spectrum, when CRISPR TP53 codependency scores were checked, highest *USP28* scores were found in cancer cell lines originating from BLCA, STAD, BRCA, LIHC, LIHC and LUAD (Fig. 2E). The leading codependency score was found in BLCA (Effect size=0.73, p= 2.2e-08) and BLCA also had the most significant value when comparing deletions to neutral copy numbers *TP53* phenocopy score (p=4.2e-06, Supp Fig. 3G). LUAD had the second most significant codependency p-value (p=3.78e-6), and is also highly ranked in comparison of phenocopy score between deletion *versus* neutral *USP28* CNV tumors (Fig. 3F). We found a positive association between *USP28* CRISPR codependency score and the effect of *USP28* deletions in *TP53* phenocopying score across cancer types (Supp Fig. 3G). Of note, that the “oncogene-tumor suppressor” dichotomy of *USP28* was reported (43), which might imply that *USP28* amplification could also result in a *TP53* phenocopy in certain contexts. However, our analysis did not support this in the majority of cancer types: out of 14 cancer types, only 3 of them had a stronger *TP53* phenocopy score in *USP28*-amplified samples than in *USP28*-deleted samples (Fig. 2E); this was the case for none of the primary cancer types for *USP28* phenocopying (BLCA, STAD, LIHC, BRCA and LUAD).

**Figure 3.**
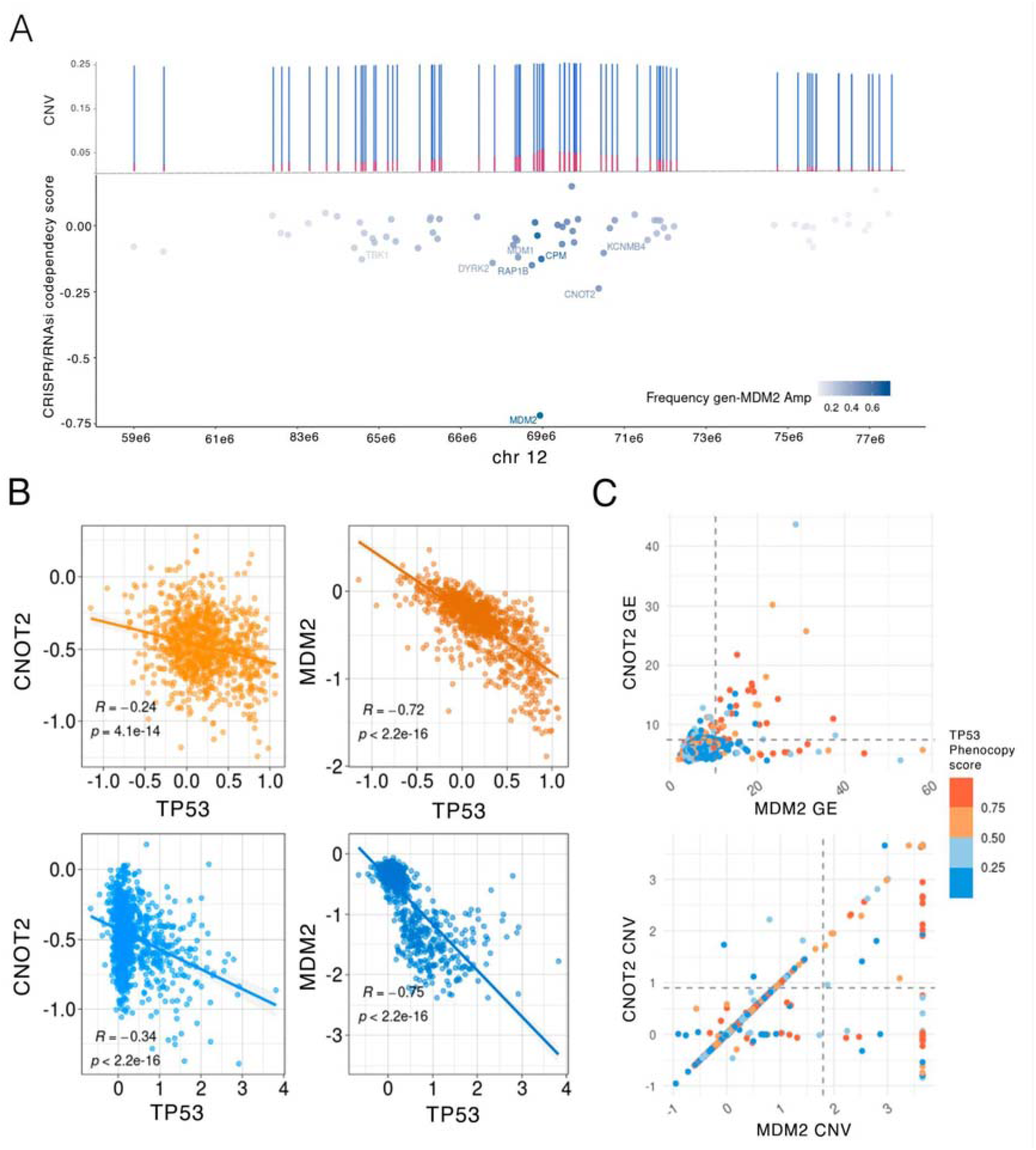
*MDM2-CNOT2* co-amplifications are associated with *TP53*-loss phenocopy score. **A.** Top: CNV of MDM2 gene neighborhoods (20Mb segment). Y axis represents the percentage of GISTIC CNV gain states +1 (blue) and +2 (red), compared to neutral CNV state (0). Bottom: CRISPR *TP53*-codependency scores (y axis) shown by gene on chromosome 12 (x axis). Genes labeled have a codependency score <-0.1, suggesting *TP53* phenocopying effects. Color shows the frequency of CNV amplification of each gene, together with *MDM2* amplifications. **B.** Co-dependency source data. CRISPR and RNAi fitness effect scores for phenocopying gene *MDM2* and candidate gene *CNOT2* (y axis), and fitness effect scores for *TP53* in the genetic screens (x axis). Top plots represent RNAi screening data and bottom plots CRISPR screening data. **C.** Association between *MDM2* and *CNOT2* gene expression (GE, top) and CNV status (bottom). Each dot represents a tumor sample, coloured based on the *TP53*-loss phenocopy score provided by the classifier. Dashed lines represent the 97th quantile across genes, for each data type.

Taken together, these results suggest that *USP28* deletion is a novel *TP53* phenocopy that commonly affects major cancer types such as breast cancer (6.2% of total breast tumors, not counting known phenocopying events and *TP53* deletions) and also bladder, lung, liver and stomach cancer (7.6 %, 7.0%, 3.8% and 2.9% cases).

### Multiple neighboring genes in a CNA segment can contribute to a *TP53* loss state

Some of the top hits found in our combined testing approach were near to known *TP53* loss phenocopying genes such as *MDM2*. We thus hypothesized that there exist cases of ‘collaboration’ of neighboring genes, affected by a single copy-number alteration, which may bear upon the *TP53* loss phenotype. This would represent a special case of epistasis between two genes, caused by a single alteration that affects both genes. Our data suggests that the *CNOT2* gene, residing in the *MDM2* segment in the chromosome 12q15, is likely an example of this relationship.

In particular, in our data, *MDM2* was frequently co-amplified with *CNOT2*, in 72% of the cases of *MDM2* amplification (Supp Fig. 4A, check by cancer type at Supp Fig. 4B). Data from CRISPR and RNAi screening experiments can help resolve associations from genomic analysis, where effects of neighboring genes are in genetic linkage (here implying being jointly affected by CNA). No other gene in that neighborhood that was amplified together with *MDM2* had as high CRISPR codependency scores as *CNOT2* (effect size=−0.24, p=4.1e-14, Fig. 3A, B); next best gene in the 20Mb neighborhood is *CDK4* with effect size=−0.16, p=3e-7. However, *CDK4* is co-amplified with *MDM2* in only 20% of the cases (Fig. 3A). *CNOT2*-only amplifications (i.e. without concurrent *MDM2* CNA) do not significantly associate with *TP53* phenocopy score (Pearson’s *TP53* phenocopy score vs *CNOT2* CNV p=0.45, effect size=−0.83, Supp Fig. 4C). More interestingly, *MDM2* CNV was not found to be associated with our *TP53* phenocopy score when *MDM2*-only amplified without *CNOT2* (Pearson’s *TP53* phenocopy score vs *MDM2* CNV p=0.57, effect size=0.09, Supp Fig. 4C). On the other hand, *MDM2-CNOT2* co-amplifications were significantly associated with a *TP53* deficiency transcriptomic score in tumors (Pearson’s correlation *TP53* phenocopy score vs *MDM2* CNV p=2e-05, effect size=0.41, Supp Fig. 4C).

This genomic evidence we provide here is supported by recent experimental studies, indicating a role for *CNOT2* in p53-dependent apoptosis, and suggesting therapeutic potential of *CNOT2* suppression (see Supplementary Text S1 for a summary and references). As supporting evidence, we considered fitness effects of CNOT2 k.o. by CRISPR in various subsets of cell lines. The MDM2-gain but CNOT2-neutral genetic backgrounds had more modest fitness effects of CNOT2 k.o. (median=−0.37) than the CNOT2-gain but MDM2-neutral genetic backgrounds (median=−0.62; p=0.072 by Mann-Whitney test, Supplementary Fig. 4D. Consistently, the *CNOT2* k.o. by CRISPR had stronger fitness effects (median=−0.55) in the TP53 *wild-type* backgrounds than in TP53-mutant background cell lines (median=−0.45, p=0.0091 by Mann-Whitney test). In other words, fitness effects of *CNOT2* disruption by CRISPR are conditional upon *MDM2* alterations and TP53 *alterations*, implicating *CNOT2* in a genetic interaction with the two other genes.

We hypothesized that this role of *CNOT2* in boosting the TP53-phenocopying effect of *MDM2* amplification may be variable across tissues. Our data suggests that in some cancer types *TP53* functional loss seems to rely on amplifications of both genes together, rather than solely *MDM2*, but not all (Supplementary Text 2). This suggests a model where the *MDM2-CNOT2* coamplification enhances the *TP53* loss effect via a genetic interaction, and of *MDM2* alone but not *CNOT2* alone able to generate a *TP53* functional loss phenotype. Gene expression profiles match this observation seen in CNA: having a *MDM2* and *CNOT2* co-overexpression (over the 97th percentile; n=40) implies a high mean *TP53* phenocopy score (above the 84th percentile, mean phenocopy score *MDM2_CNOT2=0.65*, Fig. 3C, Supp Fig. 4F), however less so for a *MDM2*-only overexpression (76th percentile; mean *MDM2* only=0.46, Fig 3 C, Supp Fig. 4F), and, expectedly, even less so for a *CNOT2-only* overexpression (73th percentile; mean phenocopy score CNOT2 only=0.41).

This principle might extend beyond the MDM2-CNOT2 pair. For instance, the *MSI2* gene, another highly prioritized hit in our combined test (Supp Fig. 4 G, H, I), resides near the known phenocopying gene *PPM1D* and thus has the potential to boost the effects of the linked amplification of the *PPM1D* gene to cause a *TP53* deficient state. Considered jointly, these findings suggest the possibility of *TP53*-loss like phenotype being a result of multiple phenocopying events generated by a single segmental CNA.

### Detecting *TP53* loss phenocopies in cancer cell line panels

It is well known that *TP53* mutations associate with overall poorer drug response in tumors (44–46), consistent with a lower ability of *TP53* deficient cells to trigger cell cycle arrest and/or apoptosis response(47–51). We hypothesized that, in addition to conferring a generalized drug resistance, the *TP53* function loss may also modulate the association between certain drugs and their target genes. In other words, we asked whether in *TP53 wild-type* cancer cells, for instance, amplification in a particular oncogene predicts sensitivity to a particular drug, while in *TP53* mutant cells the same amplification does not associate with sensitivity. Cancer cell line screening panels (52, 53) are a convenient system to test this hypothesis, because many drugs were tested systematically across both *TP53* wild-type and mutant cells of multiple cancer types. Considering *TP53* function loss via phenocopy should afford additional statistical power and clarify the associations discovered; otherwise, some effectively *TP53* null cells would be erroneously considered wild-types during association testing, making it more difficult to identify associations.

First, we aimed to generalize our tumor *TP53* phenotype classifier to cancer cell lines. Because cell lines exhibit strong global (i.e. affecting many genes) shifts in gene expression patterns, compared to their tumor tissue of origin, we applied an adjustment methodology as in our recent work (54), using the COMBAT tool (55).

Upon adjusting gene expression data from cell lines in the CCLE and GDSC panels to make it comparable with TCGA tumor data (see Methods), we applied the *TP53* classifier and obtained ranked scores. Reassuringly, the classifier assigned a significantly higher *TP53* phenotype score to *TP53* mutated cell lines (mean *TP53*_wt=0.43, *TP53* _mut=0.83, p=1.1e-49 t-test), therefore cell line data served as an independent validation set for the classifier. Of the 610 cell lines labeled as *TP53* mutant based on genomic sequence (see Methods), 87% were classified as *TP53*-loss phenotype (Fig. 4A), suggesting a reasonable ability of the classifier trained on TCGA tumors transcriptomes to generalize to cell line data.

**Figure 4:**
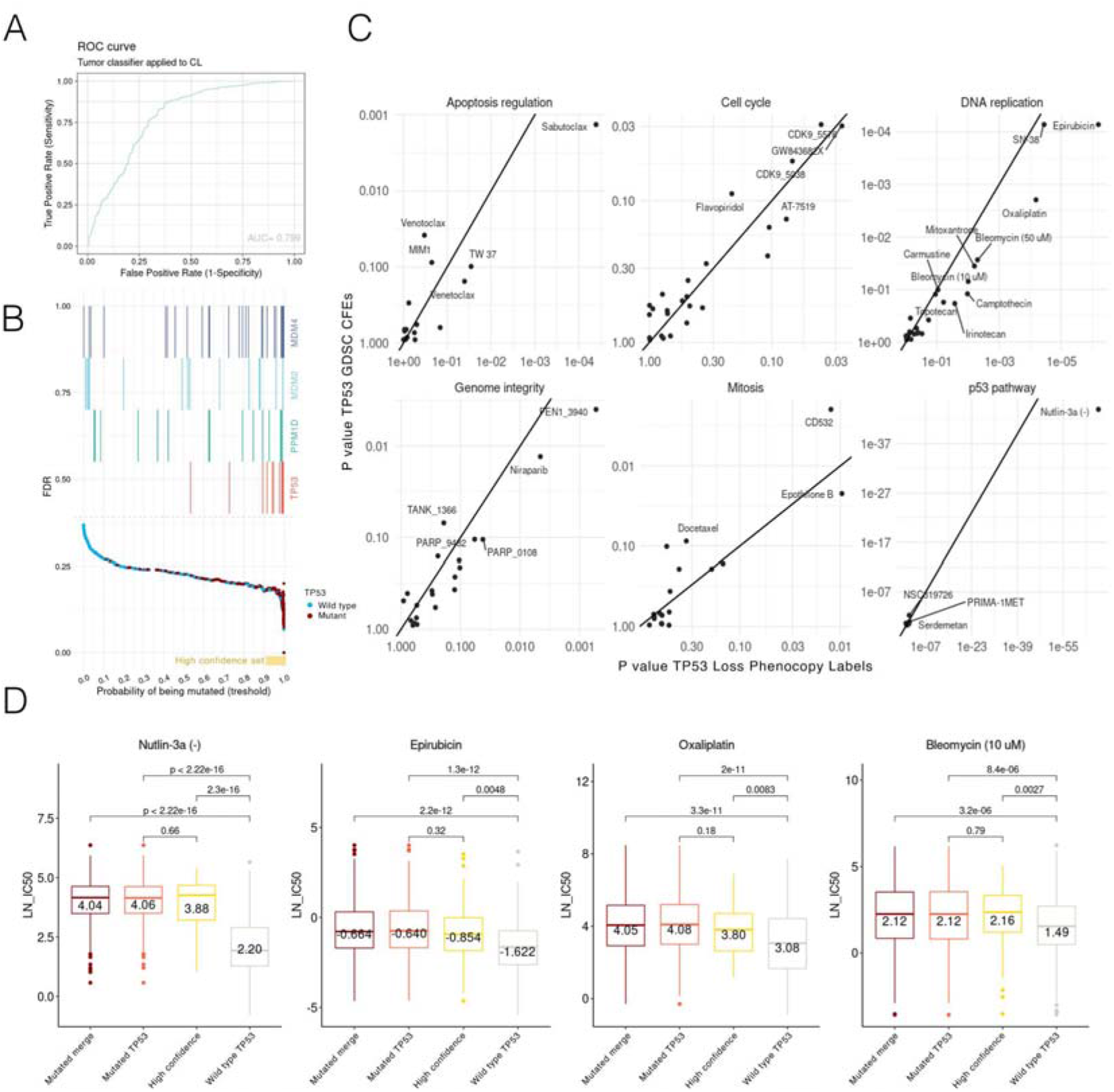
*TP53* loss phenocopy as estimated by the transcriptome score impact drug sensitivity. A. *TP53* functional status classifier, derived from tumors, is applied to cancer cell lines. Receiver operating characteristic (ROC) curve and area under curve (AUC) are shown. B. The false discovery rate (FDR) for each cell line is shown as a dot. X axis represents the phenocopy score threshold at which each cell line would be classified as *TP53* functionally deficient. Yellow horizontal bar represents the range for the high-confidence set t of TP53 phenocopying cell lines (FDR=0.18, threshold=0.93). In the top part of the plot, cell lines harboring deletions of *TP53*, and amplifications of known phenocopying genes *MDM4, MDM2* and *PPM1D* are marked. C. *TP53* status - drug sensitivity associations. Each panel represents drugs targeting genes in a given pathway. Each dot represents an association of a drug with two possible *TP53* functional status labels: X axis with the *TP53* phenocopy score and Y axis with the *TP53* mutational status (“CFE” labels by the GDSC, see Methods). P-values are from a pan-cancer regression of a given drug log IC50 versus the *TP53* status. The Y axis represents the same but using *TP53* labels according to GDSC. Associations with FDR<0.25 are labeled. D. Distributions of log IC50 values for several example drugs where *TP53* status is known to confer resistance. The X axis illustrates the different categories based on *TP53* mutated status (“Mutated *TP53’’*), wild type TP53 (“Wild type TP53”) and a high *TP53* phenocopy score (“High confidence”); the “Mutated merge” is a combination of the two. Statistical tests results comparing the groups (Mann-Whitney test, two-sided) are plotted on top. Median values are provided inside of each box.

Similarly, as in tumors, a notable fraction of cell lines were apparent false positives, labelled as *TP53* wild-type by the genomes, but classified as *TP53* deficient using the phenocopy score. We stratified these apparent false positives into a high-confidence set (“high-confidence set”); the *TP53* phenotype score of the *TP53* deleted tumor samples was used as the threshold (see Methods). The high-confidence set was composed of 76 cell lines (FDR=18%, see Methods, Fig 4 B). Only 79% of the total number of cells labelled as *TP53* wild-type genetically were also classified as *TP53* wild-type by the phenocopy score, suggesting that *TP53*-loss phenocopying events are common among cancer cell lines. In comparison, this percentage was 77% in cancer samples.

Some of the apparent false positive cell lines had a *MDM2, MDM4* or *PPM1D* amplification or a *USP28* deletion (43 out of 109, 39% of the high-confidence set). Samples harboring one of these CNA in known phenocopying genes were assigned higher scores than the rest of *TP53* wild-type cell lines (mean score=0.58 and 0.37, respectively; t-test p=5.4e-5, Supp Fig.5A). Cells harboring a *TP53* deep deletion (90th percentile of CNA scores) also had higher phenocopy scores than samples without deletion (mean score=0.78 and 0.33, respectively, t-test p=5.4e-8). 28% of the cell lines in the high-confidence harbor a *TP53* deep deletion (22 out of 76, 90th percentile of *TP53* deletion CNA). These data support that the apparent false positives are often *bona fide TP53* phenocopying events in cancer cell lines. All *TP53* phenocopy scores and cell line functional *TP53* status information is provided in Supplementary Data 4.

### Effects of *TP53* on general drug resistance are clarified by *TP53* phenocopy scores

Next, we considered the GDSC drug response distributions for various drugs, in light of the *TP53* functional status, as determined by the *TP53* mutations, or alternatively by our TP53 phenocopy scores. To identify drugs to which response is affected by *TP53* mutation status, we predicted drug response (log IC50) values of 449 GDSC drugs individually, using *TP53* status as an independent variable (see Methods).

For most of the tested drugs (105 out of 188 drugs that were significantly associated at <25% FDR, pan-cancer), the associations with *TP53* had a lower FDR when testing using *TP53* phenocopy score, over the *TP53* CFE labels (mutations which alter gene function) (Fig. 4C, effect size at Supp Fig. 5B). For the drugs that affected pathways related to *TP53* functionality, this effect of improved significance by using the phenoscore was prominent (hits FDR *TP53* phenocopy score < *TP53* CFE labels: DNA replication, 12/12 drugs, genome integrity, 8/10, p53 pathway, 3/5, Apoptosis regulation, 4/6, Cell cycle, 4/7, Supp Fig. 5C). As a negative control, randomized *TP53* labels were not significantly associated with any drug. As a positive control, the drugs known to be affected by *TP53* status such as nutlin-3a (Effect size= 1.48 vs 1.01, p= 6.7e-68 vs 1.2e-44) or bleomycin (Effect size=0.25 vs 0.16, p= 0.009 vs 0.07), exhibit a stronger association with the TP53 phenotypic score than with *TP53* CFE mutation (Fig. 4C).

We examined the IC50 drug sensitivity values of all drugs together, considering the different groups of cell lines defined by our TP53 functional status classifier (Supp Fig. 5D). Here, the mean IC50 values of our high-confidence cell lines is more similar to the *TP53* mutated cell-lines than to the *TP53* wild-type cell lines. In drugs known to be affected by *TP53* status, such as bleomycin, (Fig. 4D), IC50 values were not notably different between *TP53* mutant and the *TP53* phenocopying high-confidence cell lines. All drug associations effect size and p-value are plotted in Supplementary Figure 6 A, B. Cancer type-specific associations are shown at Supplementary Figure 6 C.

Taken together, the above analyses support the utility of the phenocopy score in identifying *TP53*-associated drug sensitivity, and also support that our tumor-derived classifier is able to generalize to cancer cell line transcriptomes to detect functional *TP53* loss phenotype.

### Associations between drug sensitivity and genetic markers is modified by functional *TP53* status

A central goal in personalized cancer medicine is to discover actionable mutations, which are used as genetic markers to decide which therapy to apply. Based on the role of *TP53* mutations in dysregulating various processes relevant to tumorigenesis, we hypothesized that various druggable cancer vulnerabilities may be conditional on *TP53* functional status. To investigate, a regression was fit to predict activity (log IC50) for each drug, from cancer type and each cancer gene mutated status (via the CFE classification, see Methods) and additionally introducing *TP53* status (either via *TP53* mutation (CFE), or via phenocopy status) as an interaction term. Comparing *TP53* phenocopy FDRs against *TP53* mutation FDR suggested that use of phenocopy score allowed to more confidently identify the drug-gene associations where *TP53* status modulates the effect size; see the comparison of FDR values (Fig. 5A), broken down by pathway that targets the drug. Out of the identified three-way associations (gene x drug x *TP53* status), 34% were found only by using the *TP53* phenocopy score, but not by the *TP53* mutation status (Fig. 5A), while for comparison only 15% are uniquely identified by TP53 mutation status. We provide a tally of all gene-drug associations that were conditional upon *TP53* in Supp Fig. 7A and a by-gene comparison of associations identified with *TP53* phenocopy score labels, versus those identified by *TP53* mutational status, in Supp Fig. 7B.

**Figure 5.**
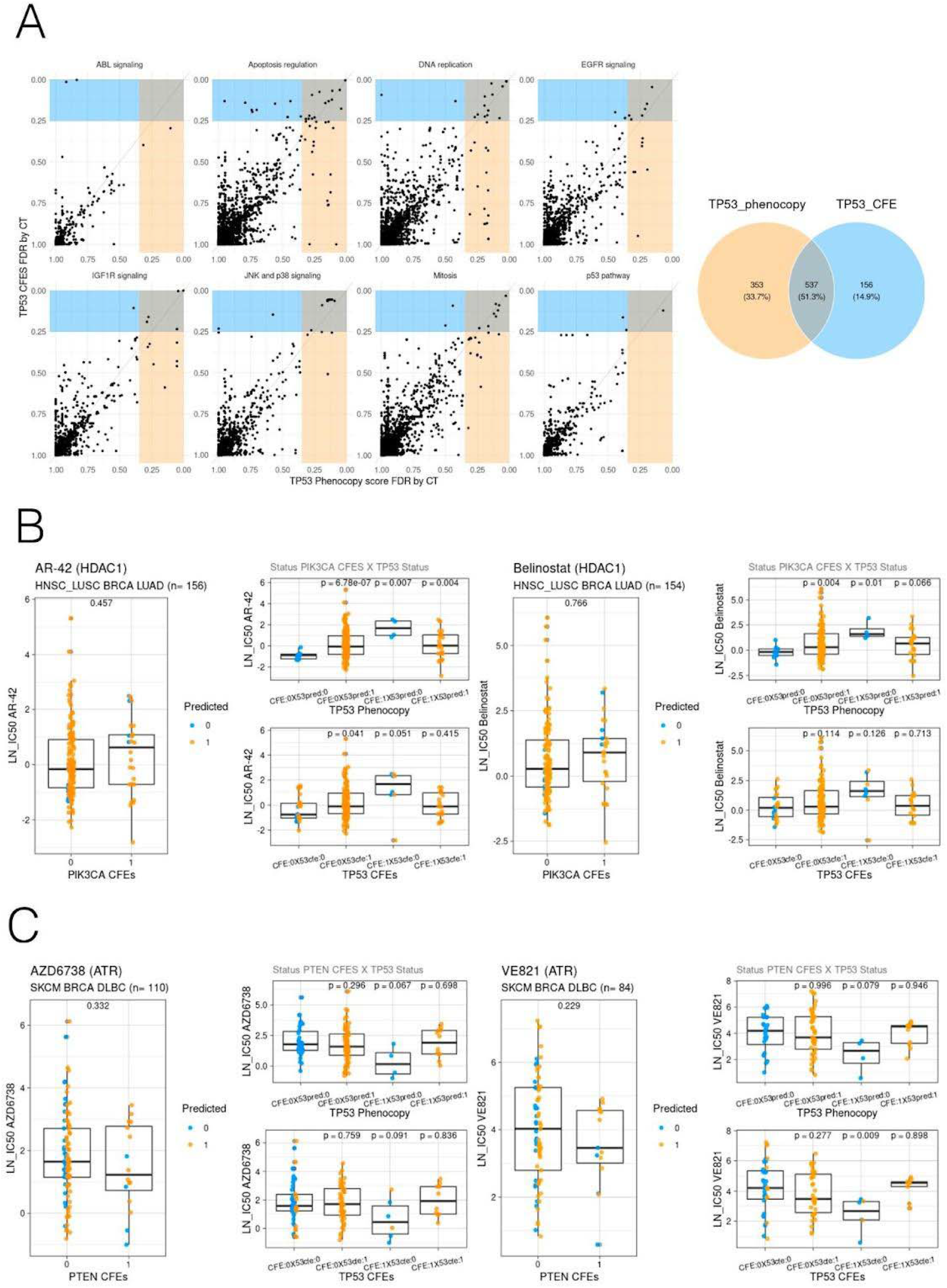
Associations between drug response and genetic markers are commonly affected by *TP53* functional status. **A.** Associations of mutations in various genes with antitumour drug sensitivity, controlling for *TP53* status. Each panel represents a pathway targeted by drugs, and each dot represents a gene - drug - cancer type combination. Associations are conditioned on *TP53* status by including an interaction term in the regression, where the Y axis shows associations using *TP53* mutational status using GDSC labels (*TP53* CFEs), while the X axis represents the same using *TP53* phenocopy score-based labels. Yellow-shaded area contains associations with FDR<0.25 for *TP53* phenocopy labels, and blue-shaded area shows the same for *TP53* CFEs. Total counts of associations in shaded areas are shown in the Venn diagram. **B.** Association of *PIK3CA* mutation status with HDAC1 targeting drugs (AR-42 and CAY10603), after controlling for *TP53* status. Large plots show the association without stratification by *TP53* labels. “CFE” denotes mutated (1) or wild-type (0) *PIK3CA* state. An association p-value is shown on top of each box by Mann-Whitney u-test. Each dot is a tumor sample belonging to one of the cancer types listed above the panel. Dots are colored according to *TP53* phenocopy score labels. Small panels represent the same association but upon stratification by TP53 status. Top row, stratification using TP53 phenocopy score labels; bottom row, using TP53 CFEs (“cancer functional events”, functional mutation status, see Methods). The X axis represents tumor samples stratified by both the PIK3CA and TP53 status. PIK3CA CFEs groups refer to PIK3CA stratification (1=mut, 0=w.t) ignoring TP53 status. Labels are as follows: “CFE:(1/0)X53pred:(1/0)” refers to stratification of PIK3CA (CFE i.e. driver mutation status) using TP53 phenocopy labels (53pred). “Last CFE:(1/0)X53cfe:(1/0)” refers to stratification of PIK3CA (CFE) using TP53 mutation labels (53cfe). “CFE:(1/0)X53pred:(1/0)” refers to stratification of PIK3CA (CFE) using TP53 phenocopy labels (53pred). Lastly, “CFE:(1/0)X53cfe:(1/0)” refers to stratification of PIK3CA (CFE) using TP53 mutation labels (53cfe) **C.** Association of *PTEN* mutation status with ATR targeting drugs (AZD6738 and VE821), after controlling for *TP53* status. Organization of the plots matches Fig. 5B, C.

Next, we aimed to select the more robust associations. To this end, we applied the “two-way” testing approach to identify replicated drug-marker links (56). In this test, it is enforced that the drug-gene association replicates across two or more drugs that share the same target gene or pathway. These were tested separately for specific cancer types, comparing *TP53*-deficient versus wild-type cells. Here, this “two-way” test (56) was further modified to be able to detect interactions with a third factor, the *TP53* functional status. As an additional criterion ensuring confidence of associations, only the hits that appear in more than one cancer type were taken into consideration (as a trade-off, this will cause highly tissue-specific associations to be missed). Stratifying by *TP53* functional status, we identified a number of drug-gene CFE associations that were not significant when ignoring the *TP53* status (60 % of total, <25% FDR, Supp Fig. 7 C). This corresponds to a total of 2303 associations of a drug to specific gene mutational status by cancer type (total number of tests ignoring *TP53*=486417 versus n=402945 controlling for *TP53* status, Supp Fig. 7D). 133 associations were found in both approaches, but revealed a lower FDR when considering *TP53* stratification (mean FDR=15% versus =19% if not stratifying=5e-08); all associations from the “two-way” replication test are listed in Supplementary Data 5.

### Sensitizing effects of driver mutations on HDAC and ATR inhibitors are modulated by *TP53*

Several studies suggested a role of the drug AR-42 (a HDAC1 inhibitor) in prolonging p53 life and triggering apoptosis (57, 58). We hypothesized that, if p53 activity is impaired, this effect of HDAC inhibitors should be reduced. Interestingly, our testing reveals that mutations in the *PIK3CA* oncogene are associated with sensitivity to HDAC1 inhibition in a manner conditional upon *TP53* mutation. In other words, when *TP53* is functional, the resistance to HDAC1 inhibitor AR-42 due to *PIK3CA* mutation is higher than when *TP53* is mutant or otherwise inactivated as indicated by phenocopy score (*TP53* wild-type A PIK3CA_mut regression coefficient test p=0.005, Cohen’s d=1.3, *TP53* mutant PIK3CA regression coefficient test p=0.08, Cohen d=−0.38, Fig. 5B). We would not retrieve this association ignoring *TP53* status (test on regression coefficient only using *PIK3CA* mutation status p=0.67, Cohen d=−0.08). In particular, in LUAD the difference in AR-42 sensitivity (median of normalized log IC50 across cell lines) between PIK3CA mutant and wild-type is hardly evident: 0.26 versus 0.24 respectively, while in *TP53*-functional LUAD this difference is −0.43 (*PIK3CA* wild-type) versus 0.35 (*PIK3CA* wild-mutant). This response is observed across three different HDAC inhibitors and in three different cancer types. AR-42 and belinostat were found significantly associated with *PIK3CA* mutation in HNSC+LUSC (here considered jointly because of known molecular similarities of the cancer types), BRCA, and LUAD cancer types (Fig 5 B). Similarly, the AR-42 association with *PIK3CA* mutation was supported in the HDAC1-targeting drug CAY10603 (Supp Fig. 7E). Furthermore, when we analyzed an independent drug screening dataset, the PRISM screen (53), we were able to recover the same associations (Supp Fig. 7E). This example illustrates how being aware of *TP53* functional inactivation status, allows to detect drug-gene associations that may be specific to the *TP53* wild-type or to the *TP53* deficient backgrounds.

We also noted that the HDAC1i-PIK3CA mutation association (conditional upon *TP53* functional status) was only recovered when controlling for *TP53* phenocopy score, but not when using simply the *TP53* mutation status (per CFE method, see Methods) as an interaction term (Belinostat IC50-PIK3CA mutation Mann-Whitney test, in the *TP53* mutation wild-type background p=0.13, while in the *TP53* w.t. phenocopy labels background p=0.01, Fig. 5B). This example illustrates how the use of *TP53* phenocopy scores provides additional power to identify drug-gene associations, as already indicated by the comparison of FDR scores for many associations above (Fig. 5A).

Recent reports have pointed out the potential therapeutic benefit of ATR inhibitors such as VE-821 or VE-822 in PTEN-defective breast, glioma and melanoma cells (59, 60). ATR is a crucial kinase regulating DNA repair and safeguarding genome integrity. ATR inhibition in PTEN-deficient cells was associated with accumulation of DSBs, cell cycle arrest and induction of apoptosis (59, 60), thus based on these phenotypes we hypothesized that the functional status may modulate this effect. Inspecting our data supports that the ATR inhibitors VE-821, VE-822, and AZD6738 were associated with a lower fitness in *PTEN*-mutant cells of the SKCM, OV, BRCA and DLBC cancers (Fig. 5C, Supp Fig. 7F). This effect was however revealed only when *TP53* status was taken into consideration, since p53 defective cells had an increased survival that obscured this association (Fig. 5C, Supp Fig. 7F). Significance of the *TP53* interaction term was not reached in this particular example, probably as the number of cell lines with a PTEN mutation (but *TP53* wild-type) was low. Nevertheless, association of ATRi IC50 values was found to be more significant in a *TP53* wild type context than in a *TP53* deficient context. This means there was a more robust difference in cell fitness comparing PTEN-mutated to PTEN wild-type cells in a *TP53*-proficient background (*TP53* wild-type IC50-PTEN Cohen’s d=−0.41 vs *TP53* deficient AZD6738 IC50-PTEN Cohen’s d=−0.05).

Overall, above we highlighted two examples where *TP53* functional status modulates the association between HDAC1 inhibitors and *PIK3CA* mutations, and ATR inhibitors and *PTEN* mutations. There were however many other significant three-way associations involving *TP53* status, cancer driver gene mutations (CNA) and drugs (listed in Supplementary Data 5), for example the association between *PIK3R1* mutations and sensitivity to MET inhibitors (Supplementary Fig. 7 G).

To estimate the importance in considering TP53 in discovering drug associations, we considered overlap in associations recovered when *TP53* status was accounted for *versus* associations obtained when *TP53* status was ignored. Only 14% of significant associations of a given molecular target to driver gene alteration status were shared between two approaches (Supp Fig. 7 F), indicating that considering *TP53* status strongly alters the drug-gene links recovered from statistical testing of drug screens. The TP53 status-aware testing recovered a higher number of associations (n=12150 versus 7853, both at <25% FDR). We also noted this effect depended on the particular gene: Drug responses in genes such as KRAS or *TP53BP1* are well explained by gene mutational status alone, not benefitting from *TP53* stratification (Supp Fig. 7 G). Nevertheless, for most of the gene, their drug associations are often more confidently retrieved when *TP53* status was accounted for (e.g. *BRAF, HRAS, ATM, APC;* n=18 genes total). Overall, the above data suggests that *TP53* should be considered when matching drugs to cancer patients based on the driver mutations in their tumor, and that this *TP53* functional status should ideally be estimated via the phenocopying score rather than *TP53* gene mutations.

## Discussion

Disabling the master tumor suppressor gene *TP53* provides cancer cells with important advantages such as avoiding cell cycle arrest or apoptosis upon replication stress or DNA damage. Because *TP53* acts as a transcription factor controlling expression of hundreds of genes, a functional read-out of *TP53* activity can be obtained using gene expression data, both at the level of mRNA or ncRNA, or at the protein level (20–23). These scores were reported to have potential clinical relevance in predicting cancer aggressiveness/patient survival and therapy response(*22, 23, 61, 62*).

In this study, we developed a global transcriptome score of *TP53* deficiencies, and applied it to ~8,000 tumors and ~1,000 cancer cell lines, to answer three questions.

Firstly, we asked how common are the *TP53*-mutation phenocopying events across various human cancers. We estimated a 12% frequency of *TP53* loss phenocopies, compared to a 58% prevalence of *TP53* mutant tumors. In some cancer types such as BRCA and BLCA, the *TP53* phenocopies may constitute a high fraction of 19% and 16% tumor samples, respectively, suggesting that the *TP53* status of tumors should preferentially be measured via functional readout (here, transcriptome-wide signature) rather than considering only mutations. Supporting this notion, a recent study using a four-gene expression signature of *TP53* activity demonstrated that this significantly predicts patient survival across 11 cancer types, and that in the majority of those it performs better than considering *TP53* mutations (22).

Secondly, given the high prevalence of *TP53* phenocopies we observed, we asked if there exist additional genetic events that are associated with these phenocopies. We developed a method considering CNA profiles and gene expression in tumors, integrating external data from CRISPR and RNAi screens, which identified the *USP28* gene deletion as a common *TP53*-loss phenocopying event. This is relevant for at least five cancer types: BLCA, STAD, BRCA, LIHC and LUAD, and affects 2.9%-7.6% tumor samples therein. The same statistical methodology also highlighted additional genes neighbouring the known phenocopies *MDM2* and *PPM1D* -- *CNOT2* and *MSI2* respectively -- which are often co-amplified with the ‘primary’ gene in the CNA gain segment and may boost the resulting *TP53*-loss phenotype. This analysis provides an example of how molecular phenotypes (here, a transcriptional signature and fitness effects from a CRISPR screen) can be used to identify multiple causal genes in a CNA segment. Analogous genomics methodologies could be applied in future work to interrogate various recurrent CNA events observed in tumors, for which the causal gene(s) are often not known with confidence.

Thirdly, we asked if a better measurement of the *TP53* functional inactivation status may be impactful in terms of predicting response to antitumor drugs based on genetic markers. Given that *TP53* deficiencies have myriad downstream consequences on the cell, including e.g. suppression of cell cycle checkpoints, or inactivation of various DNA repair pathways (*4*) it is conceivable that the *TP53* background may affect the ability of various drugs to kill cancer cells, including drugs targeted towards a particular driver mutation. We searched for three-way interactions involving *TP53* status, each drug, and each mutated cancer driver gene, finding for instance that the *TP53* status modulates the selective activity of HDAC1-inhibitors on *PIK3CA*-mutant cells. The associations were filtered to retain those supported in multiple compounds targeting the same protein or pathway; enforcing agreement across multiple measurements may allay concerns of reproducibility in cell line screening databases (63–65). Recent work by us and others (56, 66) has used statistical methods to integrate over various screening datasets, considering drug and CRISPR genetic screens jointly, to improve reliability of drug-target association discovery. Our robustly supported set of drugtarget gene links (Supplementary Data 5) that may be modulated by *TP53* status provides a resource for follow-up work to validate the role of *TP53* functional status in modulating particular gene-drug associations.

The statistical method we employed to identify *TP53* loss phenocopying events draws on the expression levels of 217 genes. Given that the model’s predictive accuracy is high (demonstrated using cross-validation and application to an independent data set of cancer cell line transcriptomes), the errors it makes are of interest. While the apparent false-positives are often *TP53* loss phenocopies, as addressed extensively in this study, it would also be interesting to look into the apparent false negatives in future. These *TP53*-mutant tumors classified as *wild-type-*like by our transcriptome score were not considered here, because of their relatively modest number, making statistical analyses difficult. Going forward, analyses of genomes from larger cohorts of cancer patients may provide enough such examples to reveal mechanisms of reestablishing *TP53* activity in certain cancers. Conceivably, this may happen by normalizing expression of the *TP53*-downstream genes which have been dysregulated by the *TP53* mutation; understanding these events may inspire new avenues for therapy of TP53 mutant tumors.

The general approach presented here could be applied beyond *TP53* also to other sorts of phenocopying events which may occur in tumors. For instance, RAS pathway activation transcriptomic scores were proposed (20), and similarly homologous recombination repair scores based on mutational signatures (86,87) Conceivably, other important cancer pathways may be similarly addressed as well, analyzing their distribution across tumors to identify possible phenocopying events, as well as their implications to drug response prediction, as we have done here for *TP53* phenocopies.

## Materials and methods

### Data collection and preparation

#### Gene expression and Copy Number Alteration (CNA) data

We downloaded gene expression data (transcripts per million, TPM) from GDC Data Portal (74) for human tumor samples (TCGA) and from GDSC (52) and CCLE (75) for cell line samples (CL). We filtered out genes with missing values in more than 100 samples and selected the overlapping genes between cell lines and tumors. Cancer types with less than 10 samples were filtered out. Low expressed genes were removed (TPM < 1 in 90% of the samples) and applied a square-root transformation to TPM. Cancer types. Tumors with less than 10 samples were filtered out. In total, we have 12,419 features for 966 CL samples and 9149 TCGA samples. We collected CNA from GDC Data Portal (74) for TCGA samples and from DepMap (64) for CL samples.

#### Data alignment between tumors and cell lines

In order to later generalize the model to cell lines we proceed to align TCGA and CL data. For this, we applied ComBat, a batch adjustment method, to account for intrinsic differences between tumor signal and cell lines signal (55). For the alignment of TCGA and CL data, we first applied quantile normalization (normalize.quantiles function, preprocessCore R 1.48.0 package) using tumor data as reference and then applied ComBat (ComBat function, R package sva 3.32.1). Each group (TCGA, GDSC or CLLC) was treated as a different batch.

#### *TP53* status label (according to GDSC)

TCGA Pan-Cancer Atlas somatic mutation data were extracted from the MC3 Public MAF (v0.2.8) data set (76). We followed the Iorio et al. methodology (24) to determine bona fide *TP53* mutations (0:wild type, 1: mutated). We identify recurrent variants that are likely to contribute to carcinogenesis. We considered mutated variants: non-synonymous missense mutations, indels (in frame insertions and deletions and out of frame insertions and deletions), nonsense mutations and specific splice-site mutations (such as “p.X125_splice”). Samples without any of these mutations annotated were considered *TP53* wild type. Just in 5% of the cases (179 out of 3416) our labels differed from the ones provided by Iorio et al. In total, we obtained *TP53* labels for 7788 TCGA tumors.

#### *TP53* score classifiers in human tumors

We used the aligned human tumor data to train a supervised elastic (20–23) net penalized logistic regression (using cv.glmnet function with alpha = 0.5, R package glmnet 4.0-2) classifier with cyclical coordinate descendent optimization (77). The choice of Elastic net penalization aims to deal with two concerns: the large number of variables can lead to high complexity (overfitting) and the feature multicollinearity. Elastic net regressions are seen as a good trade-off that benefit from the dimensionality reduction provided by Lasso penalization while keeping as many informative variables as possible (Ridge penalization). Of note, these three regularization methods yielded similar cross-validation accuracy: Elastic net (i.e. alpha=0.5) AUC 0.960, Lasso (i.e. alpha=1) AUC 0.965, and Ridge (i.e. alpha=0) AUC 0.952, suggesting that the default alpha=0.5 in Elastic net method is a reasonable choice. The model is trained using RNAseq data (X matrix) to infer *TP53* status (Y matrix). As a reference (Y) during training we used *TP53* mutation status labels.

For the training set, we excluded the tumor samples that have an amplification (not neutral, >0, according to GISTIC CNA thresholded calls downloaded using FirebrowseR package, Analyses.CopyNumber.Genes.Thresholded function) in previously known *TP53* phenocopying genes (*MDM2, MDM4, PPM1D*) or a deep deletion of *TP53*, to prevent the model from relying too much on dosage effects of these genes, instead of the downstream response.

In addition, to control for cancer type specific signals we included cancer type as a dummy variable. To control for class imbalance, we included weights in the classifier.

The model learns a vector of gene-specific weights that better classifies *TP53* status. The score from the models determines the probability of a given tumor of being *TP53* deficient. Optimization of the penalized regression formula and further details of the classifier can be consulted at (77)

#### Assessment of the classifier and calculation of FDR score

Using 90% of the training set and 5 balanced folds (balanced based on *TP53* mutational state) we performed cross-validation. We measured the performance of the training set (folds used for training) and the testing set (10% held out). Areas under the Receiving Operating Curve (AUROC) and the Precision Recall curve (AUPRC) were calculated for both training (cross-validation) and testing sets.

FDR was calculated by sample using each sample probability score from the classifier as threshold for determining positive and negative samples FDR=false positive / (false positive + true positive). Samples harboring an amplification (GISTIC thresholded amplifications, FirebrowseR package, Analyses.CopyNumber.Genes.Thresholded function) of known phenocopying genes (*MDM2,MDM4,PPM1D*) or *TP53* deletions (GISTIC thresholded deep deletions, FirebrowseR package, Analyses.CopyNumber.Genes.Thresholded function)) were considered as true positives when calculating FDR.

In Figure 1B, density of known phenocopies was calculated using *MDM4, MDM2, PPM1D* (amplifications) and *TP53* (deletions) CNA over/under the 95/0.05 th quantile. All *TP53* Phenocopy scores (probabilities of being *TP53* dysfunctional) are provided at Data S2.

The classifier coefficients were analyzed using the GO enrichment tool ShinyGO (78). The 12419 genes from the gene expression matrix with a coefficient equal to zero were used as background. Full classifiers relevant coefficients are provided at Data S1.

The coefficients of the *TP53* model should be interpreted with care, for several reasons: some of these genes may change in expression levels via indirect association meaning they may not be directly regulated by *TP53;* the gene set may omit genes that are *bona fide TP53* targets if the information contained in them is redundant with other genes; and finally these genes may individually be only weakly associated with *TP53* status, since the method optimizes the expression markers’ collective power. Visualization was performed using Revigo (27).

#### *TP53* status detection in cell lines

Using the downloadedRNAseq from GDSC cell lines data we applied our trained tumor classifier to cell lines. As stated above, RNAseq data was square rooted, normalized and ComBat batch corrected. Cell line prediction performance was measured using as reference *TP53* COSMIC labels (79) combined with Iorio et al methodology (24) as we did in tumors. FDR was calculated again using samples harboring an amplification of known phenocopying genes (*MDM2,MDM4,PPM1D*) or *TP53* deletions as true positives.

Using the classifier scores we separate the cell lines high-confidence set (FDR<=18%) using as threshold reference GISTIC tresholded *TP53* deep deletions (−2) (threshold=0.93) (FirebrowseR package, Analyses.CopyNumber.Genes.Thresholded function). Therefore, we determine 3 sets derived from our Phenocopy score: high-confidence set (predicted *TP53* phenocopies, classified as mutant but originally labeled as wild type), *TP53* mutant (classified and labeled as mutant) and *TP53* wild type (classified and labeled as wild type). All cell line predictions are provided at Data S3.

Due to a lack of positive controls, samples that were classified as wild type being originally labeled as *TP53* mutant were not considered further. However, in the future, analyses with a higher number of cancer genomes may reveal mechanisms of reestablishing *TP53* activity in some *TP53* mutant cancers (e.g. by normalizing expression of the *TP53*-downstream genes which have been dysregulated by the *TP53* mutation).

#### Gene co-dependency with *TP53* knockout/knockdown

Following data of the 2021 Q4 release downloaded from the DepMap project website: CRISPR data from PROJECT Score (28) (“Achilles_gene_effect.csv”), combined RNAi from DEMETER2 project (29) (“D2_combined_gene_dep_scores.csv”), and the cell line metadata (“sample_info.csv”). In this data, negative scores imply cell growth inhibition and/or death following gene knockout.

CRISPR data is normalized so non-essential genes scores are close to 0. We used Pearson’s correlation to correlate the gene effect of CRISPR *TP53* knockout in every cell line to other genes’ effect. We tested 990 cell lines for our 12419 genes. This score was calculated both by pan-cancer and by cancer type.

Equally to CRISPR codependency data we correlated gene knockdown effect with *TP53* knockdown effect using Pearson’s correlation test. We tested 700 cell lines for our 12419 genes. This score was calculated both for pan-cancer and by cancer type.

#### Calculation of the prioritization score

We sought to rank possible *TP53* loss phenocopying genes testing different data: copy number variant data, gene expression data (RNAseq), RNAi codependency score and CRISPR codependency score. We used the downloaded tumor data (previously described) and our *TP53* Phenocopy score to test for differences across our 3 main *TP53* groups: *TP53* wild type (labeled and classified as wild type), *TP53* mutated (labeled and classified as mutated) and predicted *TP53* phenocopied(labeled as wild type but classified as mutated). We guessed that phenocopying genes should have a differential expression in the phenocopies group when comparing to wild type and mutated *TP53* groups individually. We tested 12419 genes (by cancer type) in the following manner (via Student’s t-test):

CNV_gene(i)_*TP53*_wt *versus* CNV_gene(i)_*TP53*_phenocopies (CNV0 test),
CNV_gene(i)_*TP53*_mut *versus* CNV_gene(i)_*TP53*_phenocopies (CNV1 test)
GE_gene(i)_*TP53*_wt *versus* GE_gene(i)_*TP53*_phenocopies (GE0 test)
GE_gene(i)_*TP53*_mut *versus* GE_gene(i)_*TP53*_phenocopies (GE1 test)
RNAi_score_gene(i) *versus* RNAi_score_TP53 (RNAi codependency score, methodology described above)
CRISPR_score_gene(i) *versus* CRISPR_score_TP53 (CRISPR codependency score, methodology described above).

3010 genes out of 12419 did not have gene expression data so GE1 and GE0 tests were omitted from the combination for those genes. We combined the p-values values from the tests by cancer type using Fisher’s method for combining p-values. For each category (CNV and GE) we only use in the combination the worst p-value (max) between CNV0 and CNV1 and GE1 and GE0 as a way of controlling. Genes in which the test direction is not coherent in CNV, GE and codependency score were dropped. A gene with a negative codependency score, as negative regulators such as *MDM2*, is expected to cause a phenocopy of *TP53* by amplification and overexpression (therefore a higher expression in the phenocopies group that *TP53* wt or mut). P-values were FDR adjusted using Benjamini-Hochberg method (p.adjust function of the stats package). We further merged each cancer type combined score into one single FDR value using Fisher’s approach. That way we obtained the final Prioritization score for each gene in a cancer-combined way. We set as reference the known phenocopies (*MDM2, MDM4, PPM1D*) FDR and CRISPR codependency score. To establish a stringent threshold for new possible phenocopying genes, we determine that the gene’s prioritization score (combined by cancer type) should have an FDR as significant as the best ranked phenocopying gene (by cancer type). Same was applied for CRISPR codependency score. The known phenocopying genes with the best score by cancer type was *MDM4* in LUAD, with an FDR of 4e-05 and a CRISPR codependency score of −0.26.

#### *TP53* wild-type and *TP53* -/- isogenic cell line screens

Mean beta scores were calculated using MAGeCK-MLE (80) for *TP53*-isogenic pair cell lines A549 (81) and two RPE1 cell lines (82, 83). Beta scores represent the effect that gene knock-out has on cell fitness.

We calculated the Z-scores (distance from the mean expressed as number of standard deviations) of either *USP28* or ATM within the distribution of their respective neighbor genes, for each dataset and *TP53* status “1Mbp neighbor genes” are genes present in Brunello (84) and Gecko v2 (85) libraries and located within a 1Mbp window surrounding either *USP28* or ATM, obtained from genecards.weizmann.ac.il

#### Drug response associations with *TP53* status

We collected GDSC (24) drug data for a total of 1000 cell lines. We used IC50 as a measure of activity of a compound against a specific cell line. If drug data was available in both GDSC1 and GDSC2 versions, GDSC1 data was selected.

We also collected each drug putative target and target pathway information from the GDSC website (https://www.cancerrxgene.org/). We filtered out NA values and transformed IC50 to log scale. We downloaded GDSC mutational Cancer Functional Events (CFEs) (24) in order to: make comparisons between *TP53* Phenocopy score and GDSC *TP53* CFEs and to test other gen status drug responses controlling for *TP53* status. Mutational CFEs consist of a GDSC curated set of cancer genes (CGs) for which the mutation pattern in whole-exome sequencing (WES) data is consistent with positive selection.

We first used drug response (IC50) values of 449 GDSC drugs to fit a pan-cancer regressions against *TP53* status using cancer type as control variable. We fit three different regressions per drug response: against *TP53* CFEs, against predicted *TP53* Phenocopy thresholded scores and against *TP53* random labels.

##### log(IC50) ~ TP53.status + cancer.type

For the *TP53* status we used the groups obtained from our Phenocopy score being the *TP53* high-confidence set (classified as mutant, labeled as *wild-type*) and *TP53* mutant set (classified as mutant, labeled as mutant) the *TP53* deficient set (*TP53*.status = 1) and *TP53* wild type (classified as *wild-type*, labeled as *wild-type*) as wild type set (*TP53*.status = 0). Due to uncertainty, we filtered out samples with a *TP53* mutation classified as *wild-type*. Cancer types with less than 3 cases for any category were filtered out. We used the *esc* R package to calculate effect size (cohens_d function). P-values of associations were FDR corrected using the Benjamini-Hochberg (“fdr”) correction of the p.adjust function (stats package).

We separate the drugs into groups according to the pathway the gene they target belong to. By pathway, we calculated the slope resulting from the comparison of the FDR Phenocopy score regression versus the FDR *TP53* CFEs. For visualization we plotted raw IC50 values of different drugs and all drugs together across the different cell line defined sets. For further analysis, we merged the cancer types that were similar: HNSC with LUSC (jointly known as HNSC_LUSC), GBM with LGG (LGG_GBM) and OV with UCEC (OV_UCEC).

#### Drug response associations of gene status controlling for *TP53* status

We collected drug screening data from the PRISM project (53) and GDSC project (52). NA values were filtered out and IC50 values were transformed to logarithmic scale. We downloaded mutation features (GDSC mutational CFEs, see above) from (24).

First, we fit a regression for each drug and gene CFE including *TP53* loss Phenocopy score and the interaction term as it follows:

##### log(IC50) ~ genCFEs+ TP53Phenocopy.status+genCFEs*TP53Phenocopy.status

For comparison, we performed the same analysis using *TP53* random and *TP53* CFEs instead of *TP53* Phenocopy.status.

We tested every gene mutational CFEs out of the 300 genes provided by GDSC. We filtered out cases with lss than 3 samples in any category (mutated:1 or wildtype:0) for *TP53* status and gen CFEs. Regressions were fitted by cancer type using *glm* package (glment 4.0-2 R package). We selected genCFEs p.value and FDR correct using the Benjamini-Hochberg (“fdr”) correction of the p.adjust function (stats R package). The coefficient of the genCFEs variable informs us about the fold change of the different variable states (mutant: 1-wildtype:0) when *TP53*Phenocopy.status is set to its reference levels (wildtype:0). We compared these scores when using *TP53* Phenocopy to *TP53* CFEs by plotting FDR values and calculating slope (Figure 5 A, Supplementary Figure 7 A).

#### Two-way association tests

To further analyze *TP53* interaction in a more stringent way we implemented a version of the “two-way association test” approach developed by Levatic et al (56). In this methodology we enforced that, for a given drug, an association between a gen feature (GDSC gen mutational CFEs) and GDSC drug response is reproduced in other drugs with the same molecular target (controlled by *TP53* status as an interaction).

For this, we curated 996 sets of two drugs with the same target (ie: Dabrafenib and AZ628, target=BRAF). For the two drugs separately, we fitted a regression comparing the GDSC drug response against gen status (GDSC mutational CFEs) controlling for *TP53* status (as stated above) by cancer type. We tested the different labels in the regression: *TP53* CFEs, *TP53* Random labels and *TP53* Phenocopy labels. We considered associations by cancer type. We calculated the two-way association score by averaging the estimates (effect size) obtained between drug 1 and drug 2. To calculate the p-value for each drug-drug combination, we shuffled the *TP53* labels and compared the obtained random estimates with the actual estimate as described in our previous work (56).

For an association to be selected, we require that it is observed in more than one cancer type (merged cancer types excluded), FDR<25% across all cancer types where the hit is observed and that the direction (value from gen CFEs variable estimate) is maintained across drugs. When selecting relevant hits we also required that each hit *TP53* interaction term variable in regression is significant (FDR<25%). This informs us of deviation from the behavior of the regression variables gen_status=1 and gen_status=0 when *TP53* is controlled as interaction. We filtered out cases with less than 3 samples in any category (mutated: 1 or wildtype:0) for *TP53* status and gen CFEs in a cancer type manner. Supported hits by this methodology are reported at Figure 6 B C, Supplementary Figure 7 C, D and E and in Supplementary Data 5.

In addition, as a validation for some hits we performed a “two-way” using PRISM data. In this case we enforced that, for a given drug, an association between a gen feature (GDSC gen mutational CFEs) and GDSC drug response is reproduced in the same drug using the PRISM dataset. The rest of the methodology was applied in the same manner (see GDSC “two-way test” above).

As control, we followed the same procedure of the two-way testing method but fitting regressions of IC50 ~ gen CFEs (without interaction term). FDR corrected p-values of gen CFEs coefficient in regressions with and without interaction term were compared. We made different types of comparisons: by gene associations (Supplementary Figure 7 B), molecular target-gen CFEs associations (different 2-sets of drugs can target the same molecular feature) and all associations (Supplementary Figure 7 A)

## Supplementary material

### Supplementary Text 1

CCR4-NOT is a transcription complex (CNOT), composed of 11 subunits, that plays an important role in multiple functions in terms of regulating translation, mRNA stability, and RNA polymerase I and II transcriptions (67,68). *CNOT2*, one of the CCR4-NOT subunits, plays a critical role in deadenylase activity and the structural integrity of the complex (69) among other functions. An increasing number of studies have suggested *CNOT2s* role in tumor progression, such as in metastasis, proliferation and angiogenesis (70, 71). *CNOT2* depletion and CCR4-NOT disruption have been linked to an apoptotic response via MID1IP1 and increased p53 activity (70, 72). *CNOT2* has been reported to be among the top 5 amplified genes in chromosome 12, together with *MDM2* (73). Appealingly its overexpression has been demonstrated in several cancer types such as pancreas, prostate, liver, urinary, ovarian and breast (71). Experiments inducing *CNOT2* overexpression led to increased p21 and p53 expression, decreased apoptosis and decreased TNF-related apoptosis-inducing ligand (TRAIL) sensitivity (72, 73).

### Supplementary Text 2

In BLCA, co-amplifications are associated with a higher *TP53* phenocopy score, and are more frequent than *MDM2*-only amplifications (21 out of 32 are co-amplifications, Supp Fig. 4E). In BRCA, we found almost exclusively *MDM2-CNOT2* co-amplifications and no *MDM2* only amplifications. In STAD co-amplifications of *MDM2* and *CNOT2* are more frequent (10 out of 13) than *MDM2* solely. Just GBM was found to rely more on *MDM2* only amplifications (8 out of 14, Supp Fig. 4E).

Only 3 tumor samples were *CNOT2* amplified but *MDM2*-non amplified (all 3 having a *TP53* phenocopy score lower than 0.5, Supp Fig. 4E). No cancer type relied on *CNOT2* only amplifications.

### Supplementary Data are attached as separate files

Supplementary Data 1 - TCGA TP53 Phenocopy scores Supplementary Data 2 - Gene coefficients

Supplementary Data 3 - USP28/ATM fitness effect

Supplementary Data 4 – Cell lines TP53 Phenocopy scores Supplementary Data 5 - Two-way associations

**Supplementary Figures 1-7 are given in a separate document.**

## Notes

### Competing Interest Statement

The authors have declared no competing interest.

