## Supplementary Figures S1-S7 for "Prevalence, causes and impact of *TP53*-loss phenocopying events in human tumors"

### Supplementary Figures 1-7

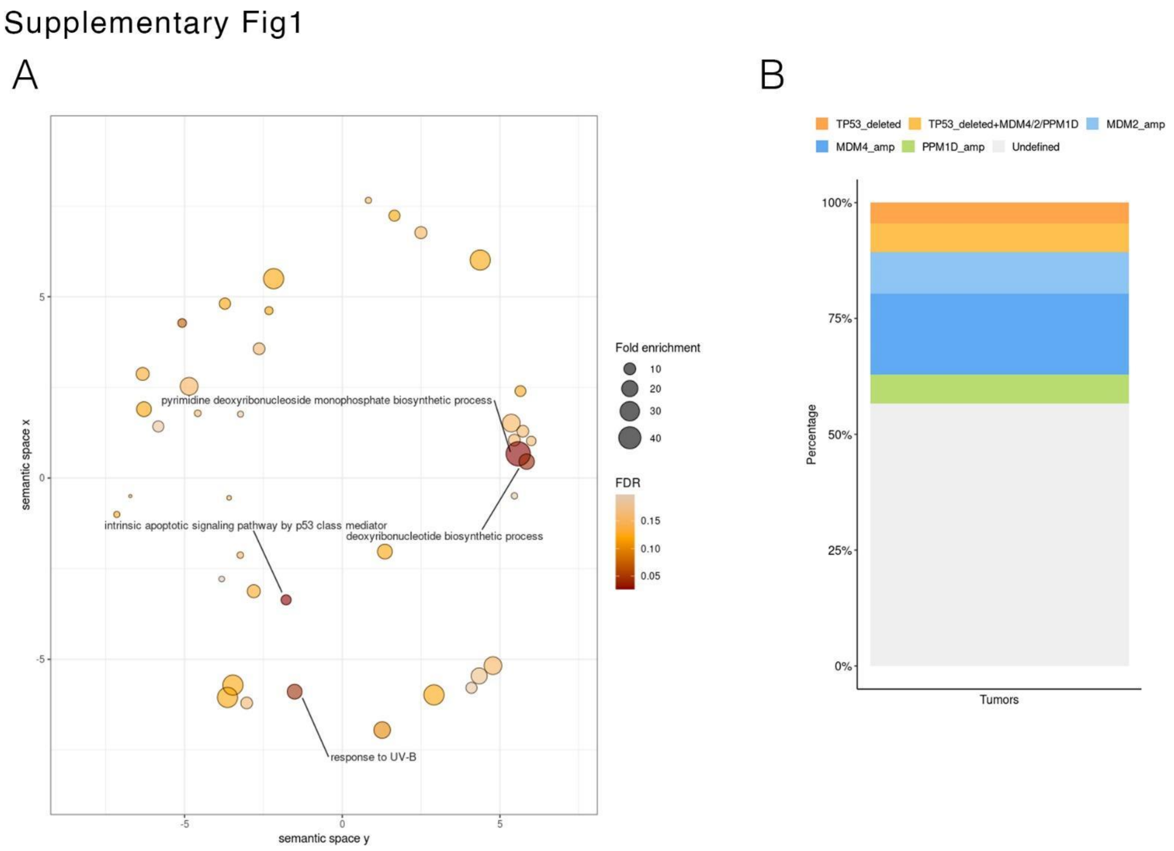


**Supplementary Figure 1. Mechanisms related to *TP53* loss phenocopying in tumors.**

A. Revigo visualization of the Gene Ontology (GO) enrichment of the *TP53* transcriptomic classifier coefficients. The axes have no intrinsic meaning, and semantically similar GO terms should remain close together in the plot (see REVIGO publication for details (27). Size indicates fold enrichment of a given GO functional category in the set of genes that had non-zero coefficients in the *TP53* classifier, while color indicates FDR of the GO enrichment.
B. Similar to Fig.1A, however the subset of *TP53* loss phenocopying tumor samples was further stratified into tumors with known phenocopying CNA events (colors), or the majority in the "Undefined" category (accounts for cases in which the cause of the phenocopy is unknown).

Supplementary Figure 2

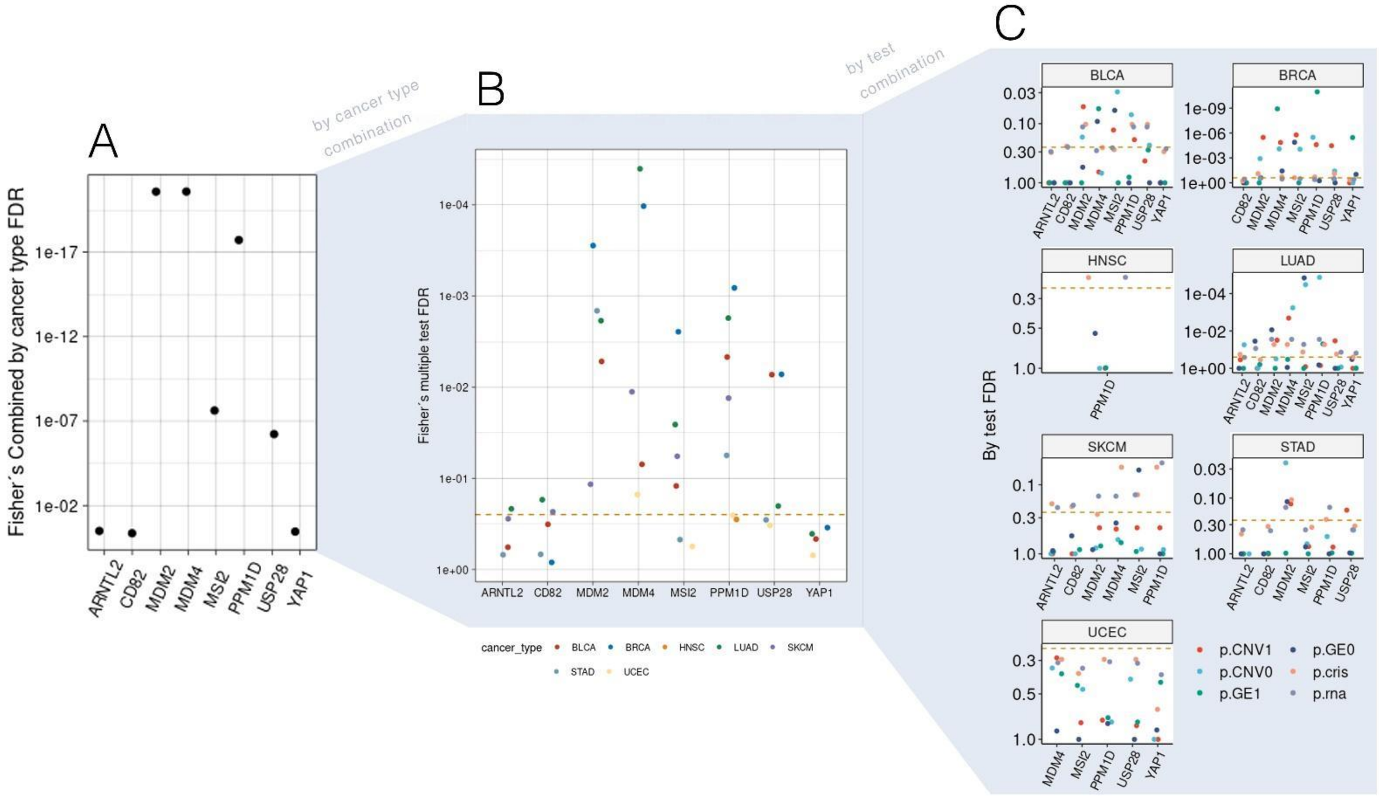


Priorization score

**Supplementary Figure 2. Prioritization score methodology**

- **A.** Prioritization score of top genes merged across cancer types. Each score combines (Fisher's method for combining p-values) six tests across four different data types (gene expression association to phenocopied samples over rest [GE], CNV association to phenocopied samples over rest phenocopy score [CNV], CRISPR *TP53* codependency, RNAi *TP53* codependency, see Methods) for each gene considered. The p-values across cancer types scores are also merged together, and final p-values are FDR corrected. The Y axis represents the final prioritization FDR of each gene. X axis represents a set of top genes, and additionally three random genes as control (*CD82, YAP1* and *ARNTL2*).
- **B.** Prioritization score p-values (FDR adjusted) stratified by cancer type. Dashed line represents 25% FDR.
- **C.** Prioritization score p-values (FDR adjusted), broken down by variable tested (colors) and by cancer type (panels). Tests compare the score of a phenocopying group of tumor samples versus the rest of tumor groups. Six different statistical tests are considered across four different data types: “pCNV1” (CNV data, versus *TP53* mutated), “pCNV0” (CNV data, versus *TP53* wild-type), “pGE1” (GE data, versus *TP53* mutated), “pGE0” (GE data, versus *TP53* wild-type), “pcris” (CRISPR codependency p-value by cancer type) and “prna” (RNAi codependency p-value by cancer type).

Supplementary Figure 3

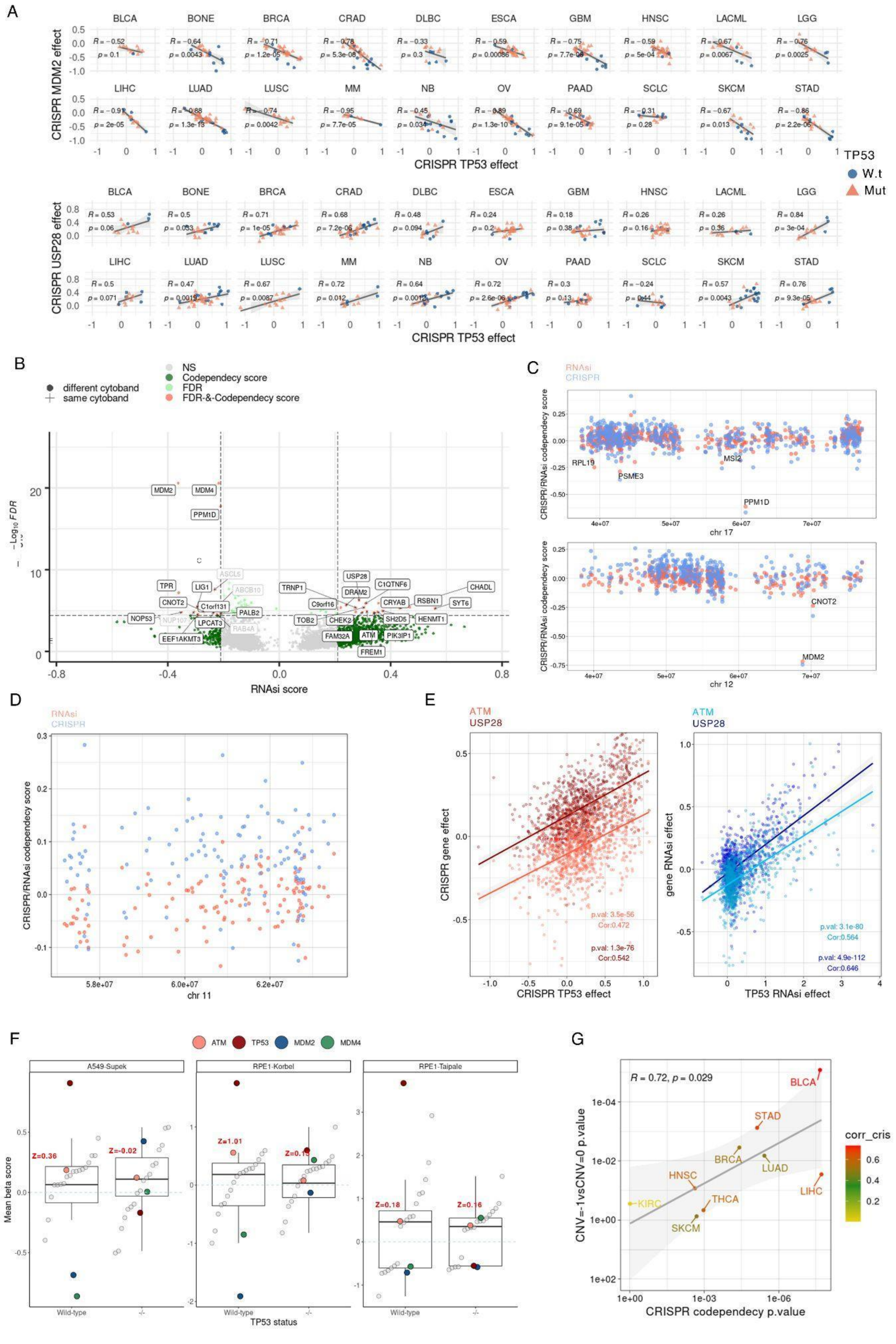

**Supplementary Figure 3. Further evidence supporting the role of individual *TP53*-loss phenocopying genes.**

- **A.** As Fig. 2A, but showing RNAi screening *TP53* codependency score on the X axis for each gene. Black colored labels are both supported by RNAi score (here) and by CRISPR score (Fig. 2A).
- **B.** CRISPR and RNAi *TP53* codependency scores of segments of chromosome 12 and 17. Genes with a score lower than -0.1 are labeled, highlighting the gene *MSI2* nearby the known phenocopying gene *PPM1D*, and the gene *CNOT2* nearby the known phenocopying gene *MDM2*.
- **C.** CRISPR and RNAi *TP53* codependency scores at chromosome 11 q12.1-q13.2 region. We found a moderate enrichment of CNV deletions in this region when comparing *TP53* loss phenocopied tumor samples against *TP53* wild-type and *TP53* mutant samples. We sought to determine whether for any genes CRISPR and RNAi *TP53* codependency scores were prominent, by analogy to *USP28* codependency scores in the q22-q23 segment.
- **D.** A comparison of the *TP53* codependency for CRISPR fitness effect across many cell lines (left) and RNAi fitness effect (right), between *USP28* and the neighboring ATM gene.
- **E.** Comparison of the mean beta score (fitness effect of CRISPR gene disruption, averaged across replicates; y-axis) of *ATM* with the mean beta scores of genes located within its 1 Mbp immediate surroundings ("1 Mbp neighbors", see Methods). Genes *TP53*, *MDM2*, and *MDM4* are also shown as a reference. x-axis bottom labels indicate the *TP53* status of the cell line. *ATM* Z-scores, comparing to the distribution of neighboring genes, are plotted in red (see Methods)
- **F.** Correlation of CRISPR screen *TP53* codependency for *USP28* (x axis), and *USP28* deletion *TP53* phenocopy scores (y axis), broken down by cancer type. Y axis represents the p-value resulting from the comparison of *USP28* neutral copy number variant *TP53* phenocopy scores versus amplified *USP28* phenocopy scores. Color represents the Pearson correlation coefficient of the CRISPR codependency.
- **G.** Comparison of TP53 Phenocopy score between USP28 amplifications (CNV=+1) and neutral (CNV=0). Y axis values represent the resulting log p-value of the statistical test comparing CNV state versus no-CNV, considering only the TP53 wild-type tumor samples. X axis represents the CRISPR codependency score broken down by cancer type (see Methods).

Supplementary Figure 4

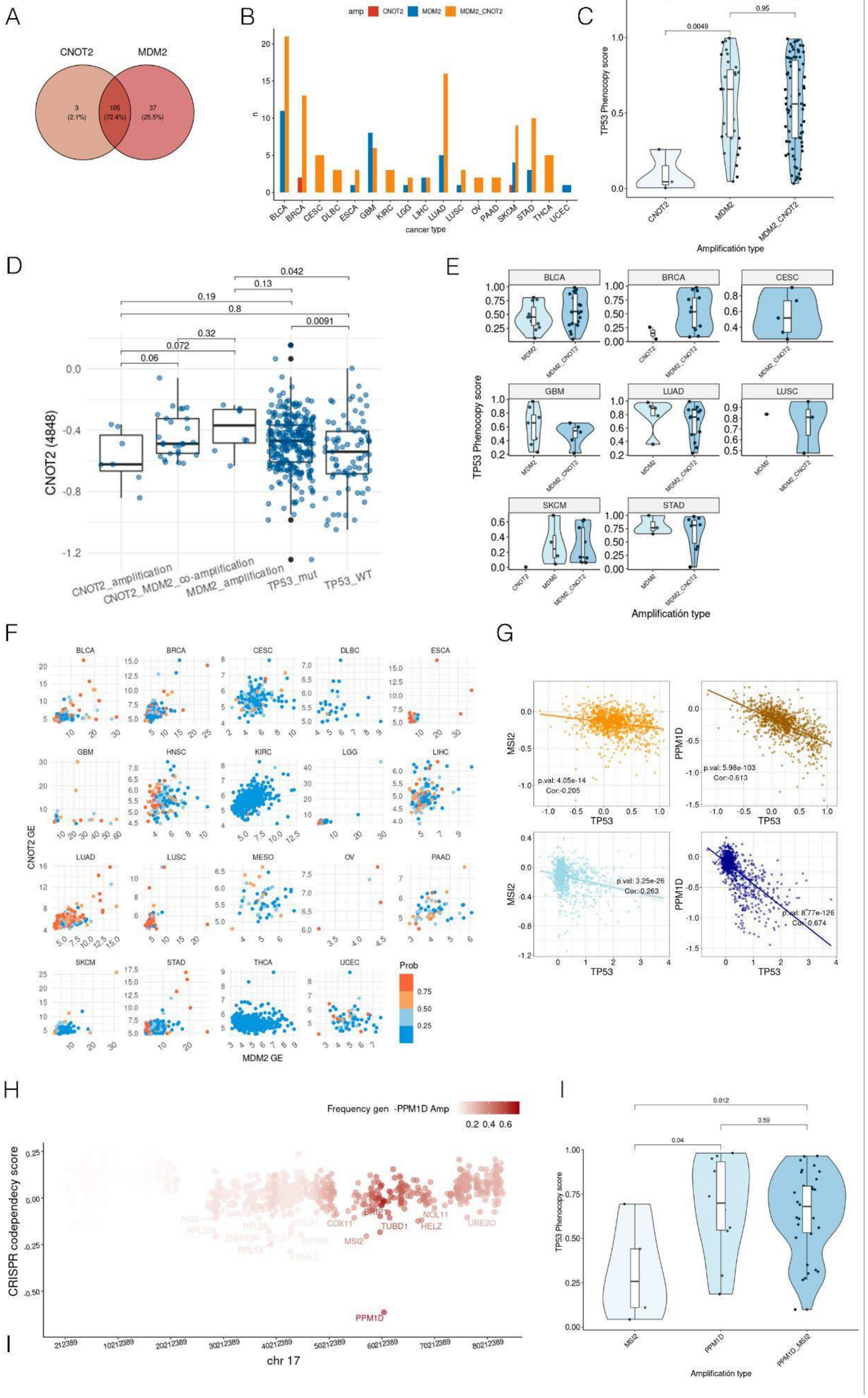

**Supplementary Figure 4. Supporting evidence for effects of *CNOT2* and *MSI2* in the amplified segments bearing the *MDM2* and *PPM1D* genes, respectively.**

- **A.** The number of tumor samples co-amplified for *MDM2* and *CNOT2,* individually for *MDM2* and individually for *CNOT2*.
- **B.** By cancer type *MDM2* - *CNOT2* CNV profiles. X axis represents different cancer types meanwhile Y axis shows the total count of events.
- **C.** pan-cancer *TP53* phenocopy scores by type of amplification. X axis represents the assigned *TP53* phenocopy score. The Y axis represents the different CNV profiles: *CNOT2* amplifications only, *MDM2* amplifications only and *CNOT2-MDM2* co-amplifications. Cancer types with less than 3 samples were removed. Each dot represents a cell line. P-value of t-test is plotted above each violin plot.
- **D.** *CNOT2* disruption fitness effects suggest genetic interactions with *MDM2* and *TP53* genes. Y axis represents the raw CRISPR score for *CNOT2* knockout (see Methods). X axis is stratified into categories based on the CCLE database CNV states of *MDM2*/*CNOT2* (thresholded at the 90% quantile) and *TP53* mutations (GDSC database CFE labels). Only *TP53 wild-type* samples were considered in every group except "TP53_mut". Each dot represents a cell line. Mann-Whitney test p-values are given above each boxplot.
- **E.** Per cancer type *TP53* phenocopy scores by type of amplification. X axis represents the assigned *TP53* phenocopy score. The X axis represents the different CNV profiles. Cancer types with less than 3 samples were removed. Each dot represents a cell line.
- **F.** Gene expression (GE) profiles by cancer type. X axis represents *MDM2* gene expression. The Y axis represents *CNOT2* gene expression. Each panel represents a cancer type. Each dot is a cell line. Colors represent the assigned *TP53* phenocopy scores
- **G.** CRISPR and RNAi raw scores for *MSI2* and *PPM1D* Top row represents RNAi codependency scores. Bottom row represents CRISPR codependency score. Right column shows *PPM1D* RNAi/CRISPR survival score in Y axis and *TP53* RNAi/CRISPR survival score in X axis. Left column shows *MSI2* RNAi/CRISPR survival score in Y axis and *TP53* RNAi/CRISPR survival score in X axis. Correlation coefficient and p-value is also pictured in each panel.
- **H.** Per-gene chromosome 17 CRISPR codependency scores. Each dot represents a chromosome 17 gene. X axis corresponds to chromosome bp position. Y axis correspond to by gene CRISPR codependency scores. Genes labeled presents a codependency score lower than -0.1. Color shows the frequency of amplification of each gene together with *PPM1D* amplifications.
- **I.** pan-cancer *TP53* phenocopy scores by type of amplification. Y axis represents the assigned *TP53* phenocopy score. The X axis represents the different CNV profiles. Cancer types with less than 3 samples were removed. Each dot represents a cell line. P-value of Fisher's T test is plotted above each violin plot.

Supplementary Figure 5

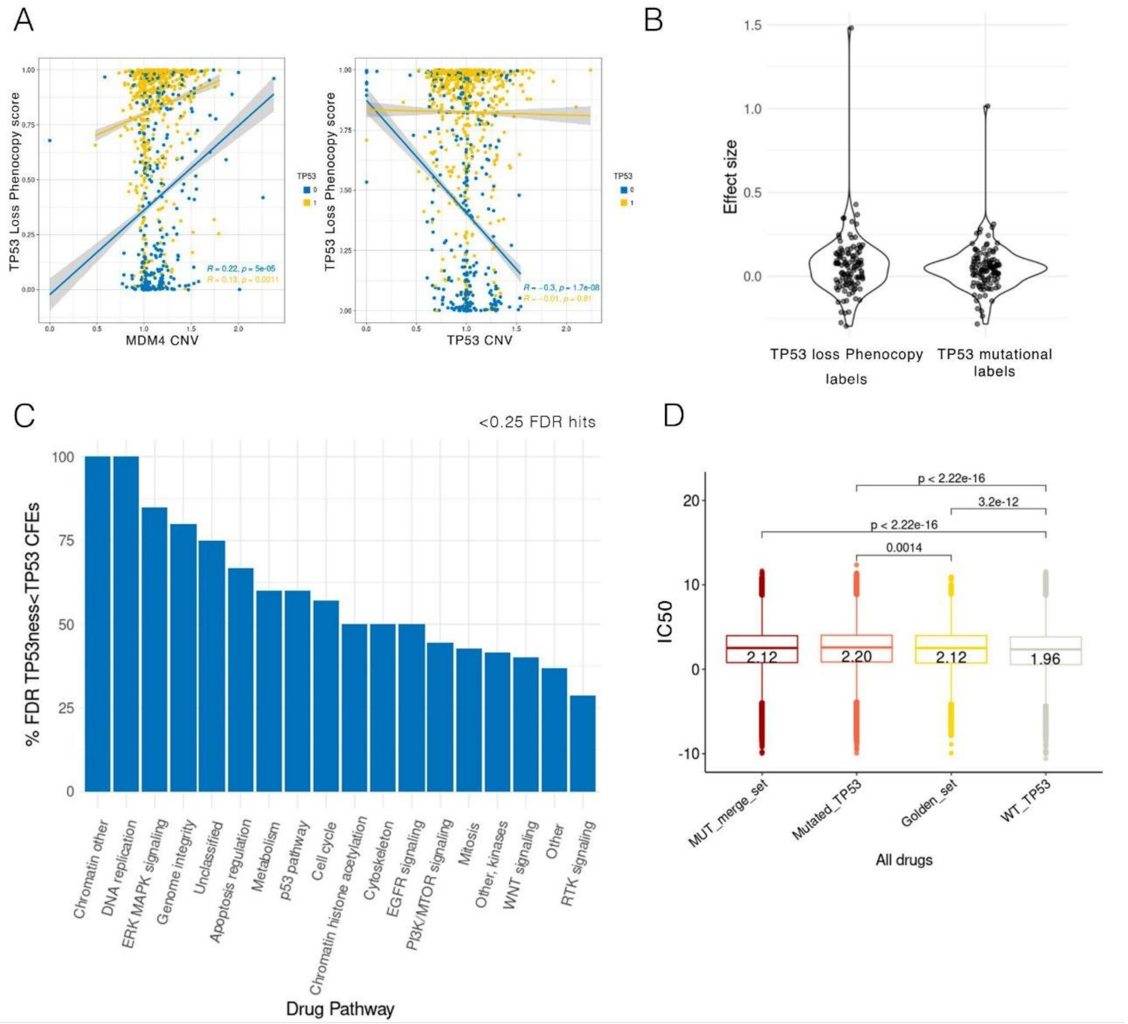


**Supplementary Figure 5. Additional data on effects of *TP53* loss and phenocopying thereof on cancer cell line drug sensitivity.**

**A.** *TP53* loss phenocopy score correlation with *MDM4* and *TP53* CNV values. Each dot represents a cell line sample colored by *TP53* status, blue for wild type and yellow for mutated. X axis represents the CNV of each given gene. Y axis represents the *TP53* loss phenocopy score obtained by applying the tumor classifier to cell lines. Two regression lines are added in each panel. Pearson's correlation R and p-value of CNV and *TP53* loss phenocopy score is also shown.

**B.** Effect size for all drugs when fitted with *TP53* status. Y axis represents the effect size resulting from fitting a regression [log IC50 ~ *TP53* status] for each drug.

**C.** Percentage of hits improved when using *TP53* phenocopy score. X axis represents the different drug categories tested. Y axis represents the frequency of hits in which using *TP53* phenocopy score improved the significance over *TP53* mutational status (CFEs, see Methods). Just hits with an FDR<25% are plotted.

**D.** Log IC50 values for all drugs in the different confidence categories obtained by our classifier. The X axis illustrates the different categories based on *TP53* mutated status ("Mutated_*TP53*") and a high *TP53* phenocopy score (high-confidence set or "Golden_set"); the "MUT_merge_set" is a combination of the two. Statistical tests results comparing the groups (Mann-Whitney test, two-sided) are plotted on top. Median values are provided inside of each box.

Supplementary Figure 6

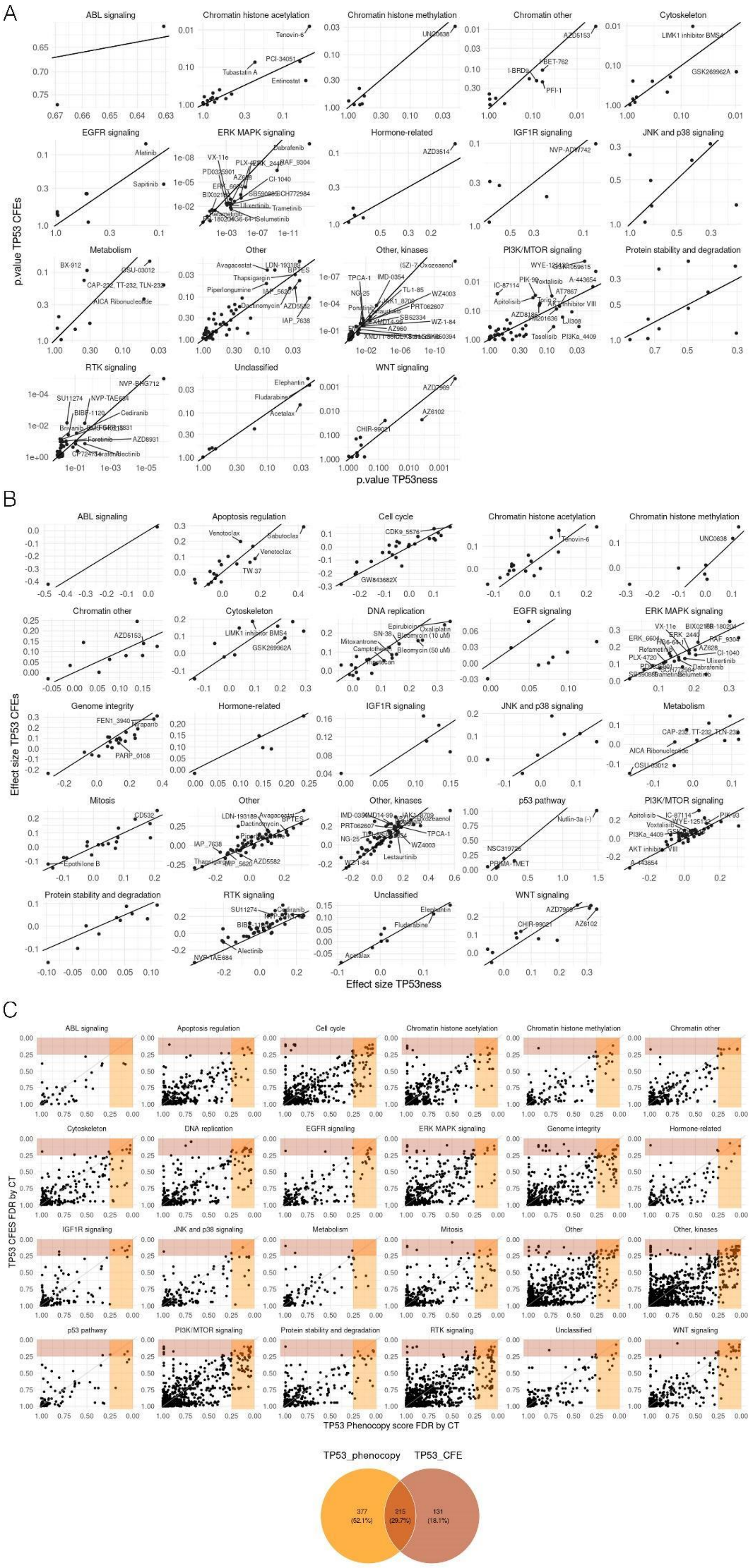

**Supplementary Figure 6. Pathway-wise effects of *TP53* mutations or phenocopies on drug sensitivity of cancer cell lines.**

**A.** Drugs tested against *TP53* status in a pan-cancer manner. Each dot represents a drug. Each panel represents a drug targeted pathway. X axis represents the p-value resultant after testing IC50 values against *TP53* phenocopy labels. Y axis represents the same but using *TP53* mutational status (CFEs). Hits under a 25% FDR are labeled. Intercept is shown as a line.

**B.** Drugs tested against *TP53* status in a pan-cancer manner. Each dot represents a drug. Each panel represents a drug targeted pathway. X axis represents the effect size (Cohen's d) resultant after testing IC50 values against *TP53* phenocopy labels. Y axis represents the same but using *TP53* mutational status (CFEs). Hits under a 25% FDR are labeled. Intercept is shown as a line.

**C.** Drugs tested against *TP53* status by cancer type. Each dot represents a drug tested in a given cancer type. Each panel represents a drug targeted pathway. X axis represents the FDR resultant after testing log IC50 values against *TP53* phenocopy labels. The Y axis represents the same but using *TP53* mutational status (CFEs). Orange colored area represents FDR<25% for *TP53* phenocopy hits, while red colores area represents the same for *TP53* CFEs. Venn diagram shows the number of hits found using each of the *TP53* labels: orange for FDR < 25% hits using *TP53* phenocopy labels and red for FDR < 25% using *TP53* mutational status (CFEs).

Supplementary Figure 7

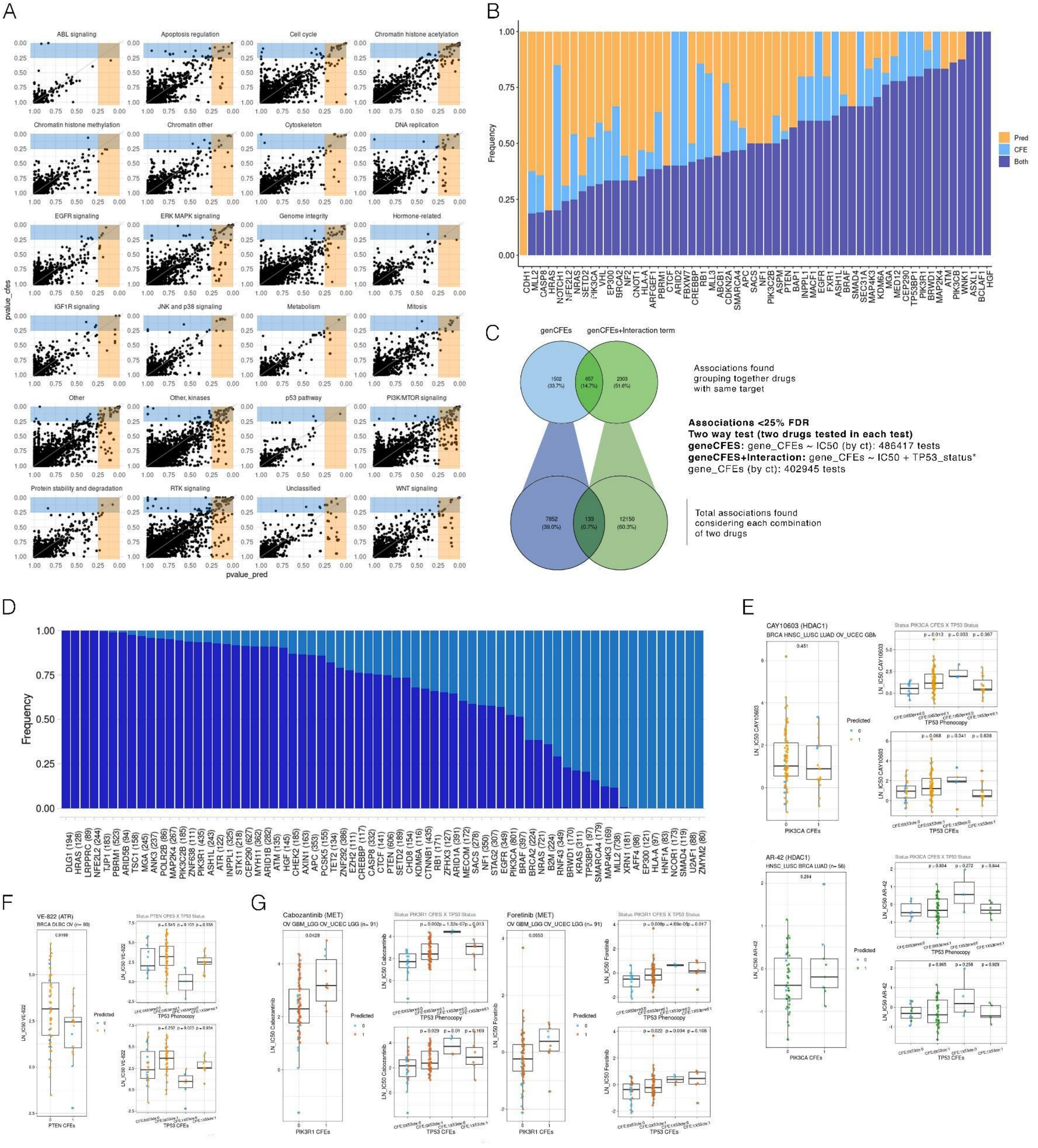


**Supplementary Figure 7. Associations between mutation status of various genes and the sensitivity to various drugs can be modulated by the *TP53* functional status.**

**A.** Associations found by pathway when testing GDSC drugs against gene CFEs using *TP53* status as interaction term. Each panel represents a pathway. Each dot represents associations of gene status with a drug targeting the given pathway, by cancer type. The Y axis represents the association´s FDR using *TP53* mutational status (CFEs). X axis represents the same using TP53 phenocopy labels. Yellow area contains associations with FDR<0.25 for *TP53* phenocopy labels. Blue area illustrates the same but for *TP53* CFEs.

**B.** TP53 phenocopy score recovery over *TP53* mutational CFEs by gene. Y axis represents the frequency of hits found by cancer type when a regression IC50 is fitted against gene CFEs using as interaction: *TP53* phenocopy score labels, *TP53* mutational status (CFE) or both. X axis represents a selection of top genes with the highest frequency of associations.

**C.** Number of hits (FDR<25%) found just using gene CFEs versus using gene CFEs plus TP53 phenocopy score status as interaction term in a regression with GDSC drugs IC50 values by cancer type. Top Venn diagram represents associations merged (hits with the same molecular target are counted together) by targeted genes - gene CFEs. Bottom Venn diagram represents all associations of drug_A-drug_B-gene CFEs.

**D.** By gene improvement of TP53 phenocopy score as interaction term. We fitted a regression by cancer type to each drug against: gene CFEs labels ignoring *TP53* and gene CFEs labels plus TP53 phenocopy score labels as interaction. The Y axis represents the frequency of hits with an FDR < 0.25. X axis represents the different genes and the number of associations found significant by cancer type (using any label). Top genes with more than 70 associations are shown.

**E.** Top (green-blue): validating the associations of AR-42 sensitivity to *PIK3CA* mutation status, when stratifying by TP53 status, using the PRISM screening data set. Bottom (Red-blue): association of another HDAC1i (Belinostat) to *PIK3CA* gene driver mutations (CFEs) when stratifying by TP53 status. Structure of the plots follow Fig. 6 B, C structure. Both associations were found by the two-way test (see Methods, hit PRISM: AR-42 & Belinostat vs *PIK3CA* CFEs, hit GDSC: CAY10603 & AR-42 vs *PIK3CA* CFEs). Just one drug of the two-way association is plotted.

**F.** VE-822 associations to PTEN driver mutations (CFEs) when considering TP53 status (two-way GDSC hit: VE-822 & VE-821 vs PTEN CFEs). Just one drug of the two-way association is plotted. Organization of the plots follow Fig. 5 B-C.

**G.** METi associations to *PIK3R1* gene CFEs when considering TP53 status. Left: association of Cabozantinib with *PIK3R1* status. Right: association of Foretinib with *PIK3R1* status. Organization is as in the Figure 6 B-C plots
